# *Bacillus velezensis*-derived muropeptide promotes growth of zebrafish via NOD2-mediated induction of IGF1 signaling

**DOI:** 10.64898/2026.03.17.712240

**Authors:** Delong Meng, Wenhao Zhou, Hui Liang, Shichang Xu, Yuanpei Zhang, Yi Wang, Yalin Yang, Zhen Zhang, Yuanyuan Yao, Qianwen Ding, Ming Li, Nan Wang, Chen Wang, Yuling Tao, Zhigang Zhou, Chao Ran

## Abstract

The role of gut microbiome in regulating vertebrate metabolism has been well-recognized. However, the effects of gut bacteria on growth have been less studied. *Bacillus* is a prevalent genus in the gut microbiota of human and animals. In this study, the effect of gut-derived *Bacillus velezensis* T23 on growth was investigated in zebrafish. *B. velezensis* T23 improved the growth of zebrafish and promoted IGF1 production in the liver and muscle, with a concomitant activation of the AKT/mTOR signaling pathway. The growth-promoting effect of *B. velezensis* T23 was not dependent on lipopeptides and polyketides. Cell wall peptidoglycan isolated from *B. velezensis* T23, as well as muramyl dipeptide (MDP), was sufficient to stimulate IGF1 signaling and growth. Further, the effect of *B. velezensis* T23 on growth and IGF1 production was abrogated in *nod2*^-/-^ zebrafish, confirming that *B. velezensis* T23 promoted growth via MDP-NOD2 signaling. Gut transcriptomic analysis indicated that *B. velezensis* T23 promoted renewal and differentiation of intestinal cells, suggesting an involvement of gut-liver axis in the effect of *B. velezensis* T23 on systemic IGF1 production. Together, our results revealed an effect of gut *Bacillus*-derived muropeptide on growth via NOD2-IGF1 signaling, and provided novel mechanistic insights in the beneficial effect of *Bacillus* spp. as probiotics.

## Introduction

The gut microbiota establishes dynamic interactions with the host, widely participating in various physiological processes such as the digestion and absorption of nutrients, energy metabolism balance, and immune disease resistance [1–3]. In recent years, the regulatory effect of gut microbiota on growth of host has been reported. The gut microbiota composition is closely related to children’s growth [4]. Under normal dietary conditions, wild mice demonstrated superior growth performance than germ-free mice, showing the contribution of gut microbiota to host growth [5]. The identity of bacterial taxa responsible for the effect of microbiota on growth of host have been revealed in some studies. The interaction between *Prevotella copri* and other microorganisms including *Bifidobacterium infantis* has a positive impact on host weight gain and intestinal metabolic function, and improves malnutrition in children [6]. Consistent with the role of gut microbiota, probiotics have been reported to promote growth in both human and animals. For instance, the supplementation of *B. infantis* can promote weight gain in malnourished children [7]. Schwarzer et al. discovered that *Lactobacillus plantarum* can improve the linear growth of malnourished mice by stimulating insulin-like growth factor 1 (IGF1) production, and the underlying mechanism was attributable to cell wall component muramyl dipeptide (MDP)-mediated intestinal nucleotide-binding oligomerization domain 2 (NOD2) stimulation [5, 8]. However, whether this mechanism can be extended to normally nutritional animals is unknown.

Among the diverse indigenous gut microorganisms, the *Bacillus* genus has attracted much attention due to its unique physiological characteristics and probiotic functions [9]. As a Gram-positive bacterium capable of forming spores, *Bacillus* is widely distributed in nature and is also an important member of the microbial communities in the intestines of human and animals [10, 11]. Among the genus, *Bacillus velezensis* is a relatively new species showing beneficial effects as probiotics, showing application prospects for both human and farmed animals [12, 13]. In particular, *B. velezensis* exhibited efficient growth-promoting effect when supplemented in the diet of farmed animals, allowing for considerable profit for the industry. However, the underlying mechanism of its action remains unclear.

IGF1 is a key signaling molecule in the growth regulation axis of vertebrates. Since the theory of the growth hormone (GH)-IGF1 axis was proposed, the crucial role of IGF-1 in mediating the growth-promoting effect, maintaining tissue homeostasis and repair has been fully elucidated [5, 14]. The biological functions of IGF1 are mainly achieved through its type I receptor (IGF1R), which phosphorylates and activates downstream protein kinase B (AKT) and mammalian target of rapamycin (mTOR) [15]. mTOR, as a core regulator of cell growth and metabolism, regulates cell growth and proliferation by promoting protein synthesis [16], and is the key executor of the IGF-1 promoting effect. The regulation of the IGF1/AKT/mTOR pathway is crucial for normal growth and development of the body [8].

Zebrafish (*Danio rerio*), as a vertebrate model, has the advantages of rapid development and a short growth cycle, enabling the observation of the growth cycle within a relatively short period, and shows unique advantages in the study of host-microbe interactions [17]. The application of germ-free zebrafish model allows for precise analysis of the physiological functions of bacterial strains in the absence of microbial interference [18]. Its high homology with mammalian genes and the ease of gene editing through CRISPR/Cas9 technology made it possible to use knockout models to verify the function of specific genes [19, 20]. Furthermore, previous studies showed that *B. velezensis*, as well as other common probiotic species, showed consistent beneficial effects in zebrafish as in other higher vertebrate animals [21, 22]. Based on this, this study used zebrafish as the model to investigate the effects of the gut *B. velezensis* on host growth and IGF1 signaling.

## Methods

### Bacterial culture

Three bacterial species, including *B. velezensis* T23 [23], *B. velezensis* T23-Δ*sfp* [23], *Plesiomonas shigelloides* CB5 [24] and *Aeromonas veronii* XMX-5 [24], were isolated from the intestine of fish in our laboratory. The cultivation of these bacteria was carried out in an oscillating incubator (Zhichu, China) with the temperature set at 30L in the Luria-Bertani medium (LB) for 24 hours.

### Zebrafish (Danio rerio) studies

All experiments and animal care procedures were approved by the Institute of Feed Research, Chinese Academy of Agricultural Sciences presided over by the China Council for Animal Care (Assurance No. 2024-AF-FRI-CAAS-001). In this experiment, we used one-month-old zebrafish samples of the AB type with similar body weights. The feeding environment was a 3-liter tank, and each group was equipped with an independent circulating water system. The zebrafish were fed twice a day for 4 weeks. The feed formula was shown in Table S1 (Each experiment had a different feed formula which was shown in the in the figure legend). Zebrafish larvae were produced from the AB strain zebrafish. The eggs were cultured in incubators for 4 dpf (post-fertilization days) until most of them hatched, and then 30 zebrafish larvae were transferred to each culture bottle containing 30 mL of filtered water. The feed formula was shown in Table S2 (Each experiment had a different feed formula which was shown in the in the figure legend). The specific details were described in the previous studies [24].

### Generation of *nod2*^-/-^ KO zebrafish

The nod2 gene of zebrafish was knocked out using the CRISPR/Cas9 technology, and a stable and heritable homozygous mutant strain was obtained. This gene consists of 11 exons. According to the targeting requirements, a target site was designed at the posterior part of exon 4. Specific guide RNA (gRNA) was designed and synthesized, and Cas9 mRNA was synthesized *in vitro*. After mixing the gRNA and Cas9 mRNA, they were introduced into zebrafish single-cell stage embryos through microinjection technology. The mutation efficiency at the target site was detected by Sanger sequencing. The F0 generation with confirmed mutations was cultured to sexual maturity and crossed with wild-type zebrafish to obtain the F1 generation. The tail fins of the F1 generation were excised to extract genomic DNA, and the target region was amplified by PCR and sequenced to screen out F1 generation individuals carrying heterozygous mutations, and to determine the mutation type. The F1 heterozygous mutant fish carrying the same mutation type were self-crossed to obtain the F2 generation. Then, the nod2 gene homozygous knockout zebrafish strain was screened out. The *nod2*^-/-^ zebrafish strain was established by the China Zebrafish Resource Center (allele: nod2ihb951/ihb951; mutation type: +5bp).

### Generation of Germ-free zebrafish larvae, as well as transferred with the gut microbiota or single bacteria

According to the previously described protocol [18, 25], germ-free (GF) zebrafish larvae were cultivated. The entire gut microbiota of the donor zebrafish was transplanted into the GF zebrafish larvae [26]. For the single strain colonization experiment, the strains *B. velezensis* T23, *P. shigelloides* CB5 and *A. veronii* XMX-5, which had been cultured to an appropriate concentration, were centrifuged at 8,000 rpm for 15 minutes to collect the bacterial cells, and then washed three times with sterile phosphate-buffered saline (PBS). Subsequently, the bacteria were resuspended to a final concentration of 10^6^ CFUs/mL and inoculated into the GF zebrafish larvae at 4 dpf. The feed was sterilized by 20 kGy γ-ray irradiation before use to ensure a sterile feeding condition.

### Zebrafish liver cells culture

The culture method of zebrafish liver cells (ZFL, ATCC CRL-2643) was as described previously [27]. When the cell confluence reaches over 80%, the culture medium is replaced, and 0.1, 1, and 10 μg/mL of MDP were added respectively for treatment. After 24 hours of treatment, the cells were collected for subsequent experiments.

### Measurement of growth performance and body indicators

Record the initial tail count (Ns) and final tail count (Ne) of each group of zebrafish, and weigh each zebrafish in each tank to obtain the initial body weight (IBW, g) and final body weight (FBW, g). At the same time, accurately record the food intake (g). After the experiment, randomly select 12 zebrafish from each group, and measure their body length (cm), fish viscera weight and carcass weight. Based on the recorded data, the following growth performance and body indicators are calculated: Survival rate (SR, %) = (Ne / Ns) × 100%; Weight gain rate (WG, %) = [(FBW – IBW) / IBW] × 100%; Feed conversion ratio (FCR) = Food intake / (FBW – IBW); Condition factor (CF, 100*g/cm³) = (FBW / body length³) × 100; Viscerosomatic index (VSI, %) = (Fish viscera weight / FBW) × 100%; Carcass ratio (CR, %) = (Carcass weight / FBW) × 100%

### Measurement of water and protein content in muscles

The moisture content in muscle tissue was determined by the direct drying method. An appropriate amount of fresh muscle sample was accurately weighed and placed in a weighed bottle that had been previously calibrated to a constant weight. The sample was then dried in a 105L oven until a constant weight was achieved after 4 hours. The moisture content was calculated based on the difference in mass before and after drying. The total protein content in the muscle tissue was determined by the Bicinchoninic acid (BCA) method (Beyotime, China). An appropriate amount of muscle tissue sample was taken, and the tissue lysis buffer was added in proportion for homogenization. After centrifugation at 12000 rpm for 5 minutes, the supernatant was collected. The operation was carried out according to the instructions of the reagent kit.

### Histological analysis of muscles and intestines

The morphology of zebrafish muscle and midgut tissues was observed and quantitatively analyzed using the hematoxylin and eosin staining (H&E) staining method. The dorsal muscle and midgut tissues of zebrafish were dissected and fixed in 4% paraformaldehyde for 24 hours. The subsequent staining and photography procedures were as described previously [28]. In the analysis of muscle tissues, the same location on each slide was selected for the field of view, and the average cross-sectional area of muscle fibers was measured and calculated using ImageJ software. In the analysis of the midgut tissue, the height of the villi and the number of goblet cells were measured using the ImageJ software.

### Detection of IGF1 content

The content of IGF1 in liver, muscle tissue and serum were detected by enzyme-linked immunosorbent assay (ELISA). After sample collection, serum samples were directly used for detection; liver and muscle tissue samples were homogenized with PBS, centrifuged to obtain the supernatant, which was stored for later use. The commercial fish IGF1 ELISA kit was used, and the operation was carried out according to the manufacturer’s instructions (Mlbio, China). The concentration of IGF1 in the samples was calculated based on the standard curve (for tissue samples, correction was required according to protein concentration or serum volume, and the unit was expressed as μg/g protein or ng/mL serum).

### Extraction of the whole cell wall and peptidoglycan components of *B. velezensis*

The extraction methods were based on previous studies [29] and was further improved. The bacterial cells were resuspended in 15 mL of lysis buffer (containing 0.01 mol/L PBS and 1 mmol/L phenylmethanesulfonyl fluoride, pH 7.4), and then ultrasonicated on ice (450 W, 5 seconds on/5 seconds off) for 30 minutes. The mixture was centrifuged at 12,000 rpm for 10 minutes to collect the precipitate. The precipitate was washed with sterile water 3-4 times to obtain the whole cell wall components. The bacterial cells were dissolved in 10% trichloroacetic acid at a ratio of 1:10 (w/v), incubated at 70°C for 1 hour, and then centrifuged at 12,000 rpm for 10 minutes to collect the precipitate; the precipitate was washed with sterile water 5-6 times, and then treated with chloroform/methanol (1:1, v/v) at 25°C for 36 hours; the precipitate was collected by centrifugation and repeatedly washed with 75% ethanol, 50% ethanol and sterile water to remove lipids; the insoluble precipitate was resuspended in a solution containing 100 μg/mL proteinase K, incubated at 55°C for 1 hour to remove proteins, and then heated at 90°C for 10 minutes to inactivate the enzyme activity; finally, the peptidoglycan was collected by centrifugation at 12,000 rpm for 20 minutes, and washed with sterile water 3-4 times to obtain peptidoglycan.

### Detection of the content of MDP

After the gut contents of zebrafish were collected and accurately weighed, they were mixed with PBS for homogenization in a ratio of mass to volume. The supernatant was then collected by centrifugation at 4°C and 12,000 rpm for 5 minutes. The supernatant samples of *B. velezensis* T23 were also collected by centrifugation at 4°C and 12,000 rpm for 5 minutes. The test was conducted using the MDP ELISA kit (CHEJETER, China). The results of the gut content samples were converted to the content per gram of gut content (ng/g), while the bacterial supernatant samples were expressed as the content per milliliter of the supernatant (pg/mL).

### Quantitative real-time PCR analysis

Total RNA was isolated from zebrafish liver, muscle, whole zebrafish larvae and the ZFL cells using TRIzol reagent (CWBIO, China). For each sample, 2μg of RNA was reverse transcribed into complementary DNA (cDNA) using the FastKing gDNA Dispelling RT SuperMix (Tiangen). The qRT-PCR assay was carried out on a Light Cycler 480 system (Roche) with the 2× SYBR Green (Tiangen). The amplification protocols and reaction mixtures are conducted as previously described [24]. Primer sequences are listed in Table S3. The ribosomal protein S11 gene (*rps11*) served as the internal reference for normalization. Relative gene expression levels were calculated using the 2^−ΔΔCT^ method.

### Morpholino knockdown

Vivo morpholino oligonucleotides (MO) synthesized by gene Tools (Philomath, OR) were used to knockdown the NOD2 gene of zebrafish larvae. The MO sequences are: NOD2-MO, 5’-TGAACTGAATCTCGCCTCATAGCCC-3’, and CK-MO, 5’-CCTCTTACCTCAGTTACAATTTATA-3’. Zebrafish larvae were immersed 50 nM NOD2 MO or standard control MO from 4 dpf. The MO was maintained throughout the subsequent experiments and the concentration remained unchanged.

### 16S rRNA sequencing of the gut microbiota

Four to six hours after the last feeding of zebrafish, the intestines were dissected and the contents were collected under sterile conditions. The total DNA of the samples was extracted using the TGuide S96 Magnetic Soil /Stool DNA Kit (Tiangen, China), and using it as the template. Amplification was performed for the V1–V9 variable region of the 16S rRNA gene using universal primers 27F (sequence: AGRGTGTGATYNTGGCTCAG) and 1492R (sequence: TASGGHTACCTTGTTASGACTT). The amplified products were purified and quantified, then mixed at equal molar concentrations, and sequenced on the PacBio Sequel II platform. The raw sequencing data was analyzed using the BMKCloud online platform (www.biocloud.net) provided by Beijing Biomarker Technologies Co., Ltd.

### Absolute quantification of *B. velezensis*

The gut contents of the zebrafish were collected under aseptic conditions. An appropriate amount of gut content samples was taken and total genomic DNA was extracted using a bacterial DNA extraction kit (Tiangen, China). The DNA concentration was detected. qPCR amplification was performed using specific primers of *B. velezensis* (the primer sequences are listed in Table S3). The Ct values of the samples were converted into copy numbers according to the standard curve. The results were expressed as the number of *B. velezensis* genes per gram of gut content (copies/g).

### Transcriptome sequencing and analysis

Extract the midgut tissue of zebrafish, extract total RNA using the TRIzol method, and test the purity and integrity of the RNA. Qualified RNA samples were used to construct a cDNA library. After quality control and inspection of the library, sequencing was performed on the Illumina NovaSeq platform. The obtained raw data were filtered for quality control and then clean reads were obtained. Used the HISAT2 software to align the clean reads to the zebrafish reference genome, and used feature counts for quantitative gene expression analysis, expressing the expression level as FPKM. Use DESeq2 to screen for differentially expressed genes (Fold Change ≥ 1.5, FDR < 0.05). Performed Gene Ontology (GO) functional enrichment and Kyoto Encyclopedia of Genes and Genomes (KEGG) pathway enrichment analysis on the differentially expressed genes. To explore the overall expression trend of the gene set, conduct gene set enrichment analysis using the Gene Set Enrichment Analysis (GSEA). Used Weighted gene co-expression network analysis (WGCNA) to construct a gene co-expression network and identify co-expression modules. The raw sequencing data was analyzed using the BMKCloud online platform (www.biocloud.net). At the same time, performed Pearson correlation analysis on key genes to calculate the correlation coefficient of expression levels between genes and reveal their co-expression patterns.

### Western blotting

The phosphorylation levels of AKT and mTOR in liver and muscle tissues were detected by Western Blot method. Appropriate amounts of tissue samples were taken, mixed with cold HBSS buffer containing protease and phosphatase inhibitors for homogenization and lysis, and the supernatant was collected after centrifugation at 4°C. The protein concentration was determined by BCA method, and the protein concentrations of each sample were adjusted to be consistent. After adding loading buffer and boiling for denaturation, equal amounts of protein were subjected to 12% SDS-PAGE gel electrophoresis separation, and then the protein was transferred to 0.22 μm PVDF membrane (Millipore, USA). The membrane was placed in a blocking solution containing 5% bovine serum albumin at room temperature for 2 hours, and the corresponding primary antibodies including anti-p-AKT (CST, 4060, 1:1000), anti-p-mTOR (MCE, HY-P80837, 1:1000), and anti-GAPDH (CST, 2118, 1:2000) were incubated at 4°C overnight. The next day, the membrane was washed thoroughly with TBST and incubated with HRP-labeled secondary antibody at room temperature for 2 hours. After washing, the membrane was exposed for imaging using Immobilon Western HRP Substrate (Merck-Millipore, Germany). The gray values of the bands were measured using image analysis software such as ImageJ, and the relative expression level of the target protein was expressed as the ratio of the gray value of the target protein to that of the internal reference protein.

### Statistical analysis

The data were statistically analyzed using GraphPad Prism 8 software. All data were presented as the mean ± SEM. For comparisons between two groups, the data were first subjected to normality test and homogeneity of variance test. Student’s t-test was employed to compare between two groups. Gut microbiota sequencing was compared by using Mann-Whitney U test. The statistical test and number of samples used are included and described in each figure legend.

## Results

### The growth promoting effect of *B. velezensis* T23 involved stimulation of the IGF1 pathway

To confirm the growth-promoting function of *B. velezensis* T23, we conducted a 4-week feeding experiment. The results showed that without affecting the SR (Fig. S1), *B. velezensis* T23 significantly increased the FBW and WG of zebrafish (Fig. 1A and B), with reduced FCR (Fig. 1C), indicating an improvement the growth performance. Balanced growth ability is an important indicator for evaluating the health of fish. In terms of physical indicators, there was no significant difference in the body length of zebrafish between the two groups (Fig. 1D), but the CF was significantly increased in the T23 group (Fig. S2). The edible part-related indicators showed that compared with the CON group, the VSI in the T23 group showed a downward trend (Fig. 1E), and the CR showed an upward trend (Fig. 1F). Moreover, muscle nutrient composition analysis revealed that muscle moisture of the T23 group was reduced (Fig. 1G), while muscle protein content was increased (Fig. 1H). Additionally, muscle histological observation results indicated that the muscle fibers of the zebrafish in the T23 group was significantly larger than that of the CON group (Fig. 1I and J).

**Figure 1.**
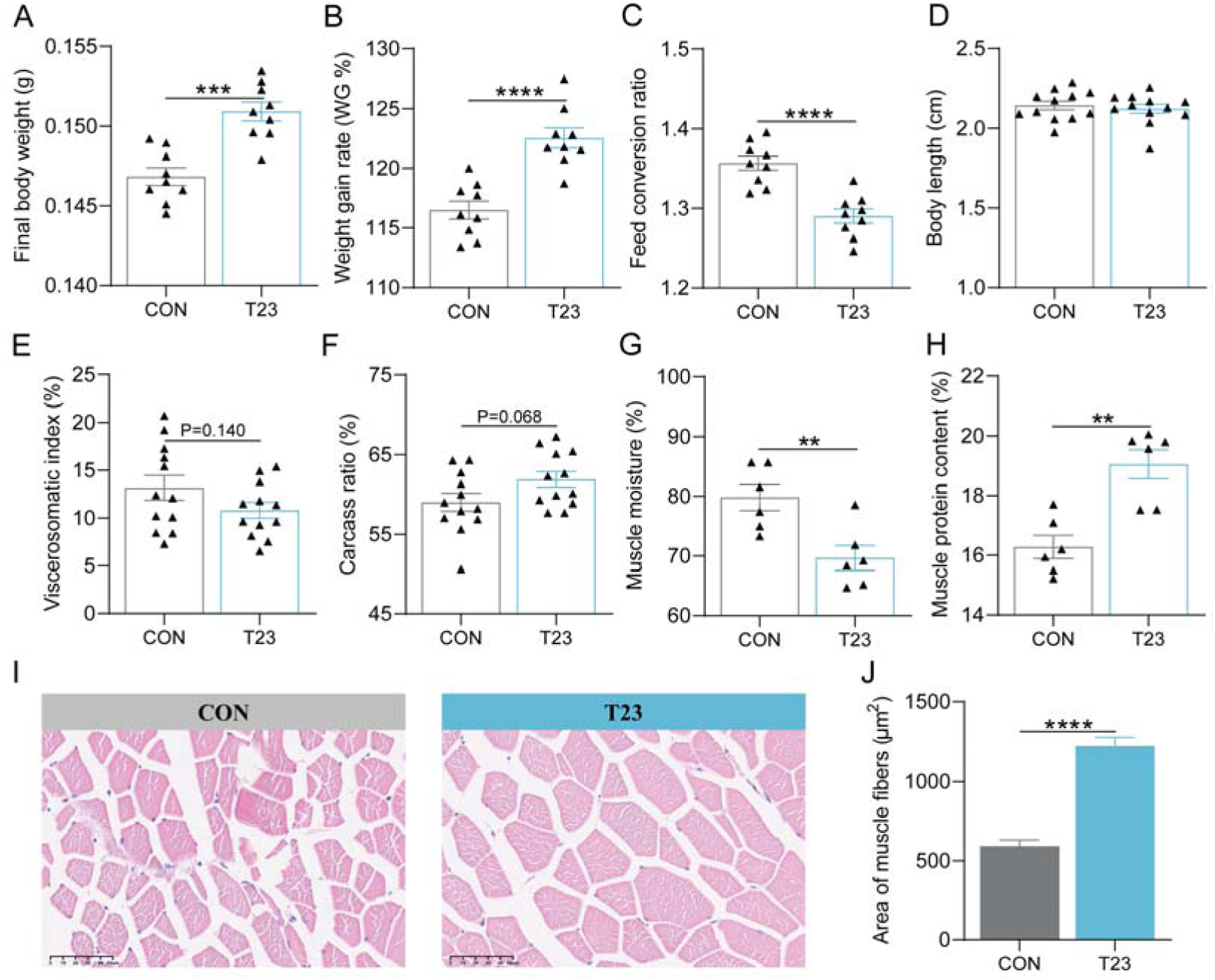
*B. velezensis* T23 improved growth and muscular protein deposition in zebrafish. One-month-old zebrafish were fed with normal diet (CON group) or diet supplemented with 10L CFUs/g *B. velezensis* T23 (T23 group) for 4 weeks. (A) Final body weight (FBW, n = 9). (B) Weight gain rate (WG, n = 9). (C) Feed conversion ratio (FCR, n = 9). (D) Body length (n = 9). (E) Viscerosomatic index (VSI, n = 9). (F) Carcass ratio (CR, n = 9). (G) Muscle moisture (n = 6). (H) Muscle protein content (n = 6). (I) Muscle H&E section (n = 3). (J) Area of muscle fibers (n = 90). Data were presented as mean ± SEM. Student’s t-test was employed to compare between two groups. Significant differences were indicated as \*\**p* < 0.01, \*\*\**p* < 0.001, \*\*\*\**p* < 0.0001.

As an endocrine determinant of growth, the liver is the main source of circulating IGF-1 [5]. *B. velezensis* T23 significantly increased the content of IGF1 in the liver and serum of zebrafish (Fig. 2A and B). Compared with the CON group, the gene expression of *igf1*, *igf1ra* and *igf1rb* in the liver of zebrafish was increased in T23 group (Fig. 2C), and the phosphorylation of AKT was significantly enhanced (Fig. 2D and E). IGF1 could also be produced by peripheral tissues (including muscles) through autocrine/paracrine mechanisms, directly promoting the growth of local tissues [5]. In muscle tissues, *B. velezensis* T23 significantly increased IGF1 content (Fig. 2F), and upregulated the gene expression of *igf1*, *igf1rb* and *ghrb* (Fig. 2G). Further analysis showed that *B. velezensis* T23 treatment significantly enhanced the phosphorylation levels of AKT and mTOR in muscle tissues (Fig. 2H-J). These results indicated that *B. velezensis* T23 activated the IGF1 signaling pathway in the liver and muscle, thereby enhancing protein synthesis mediated by AKT/mTOR and promoting muscle growth, ultimately improving overall growth performance of zebrafish.

**Figure 2.**
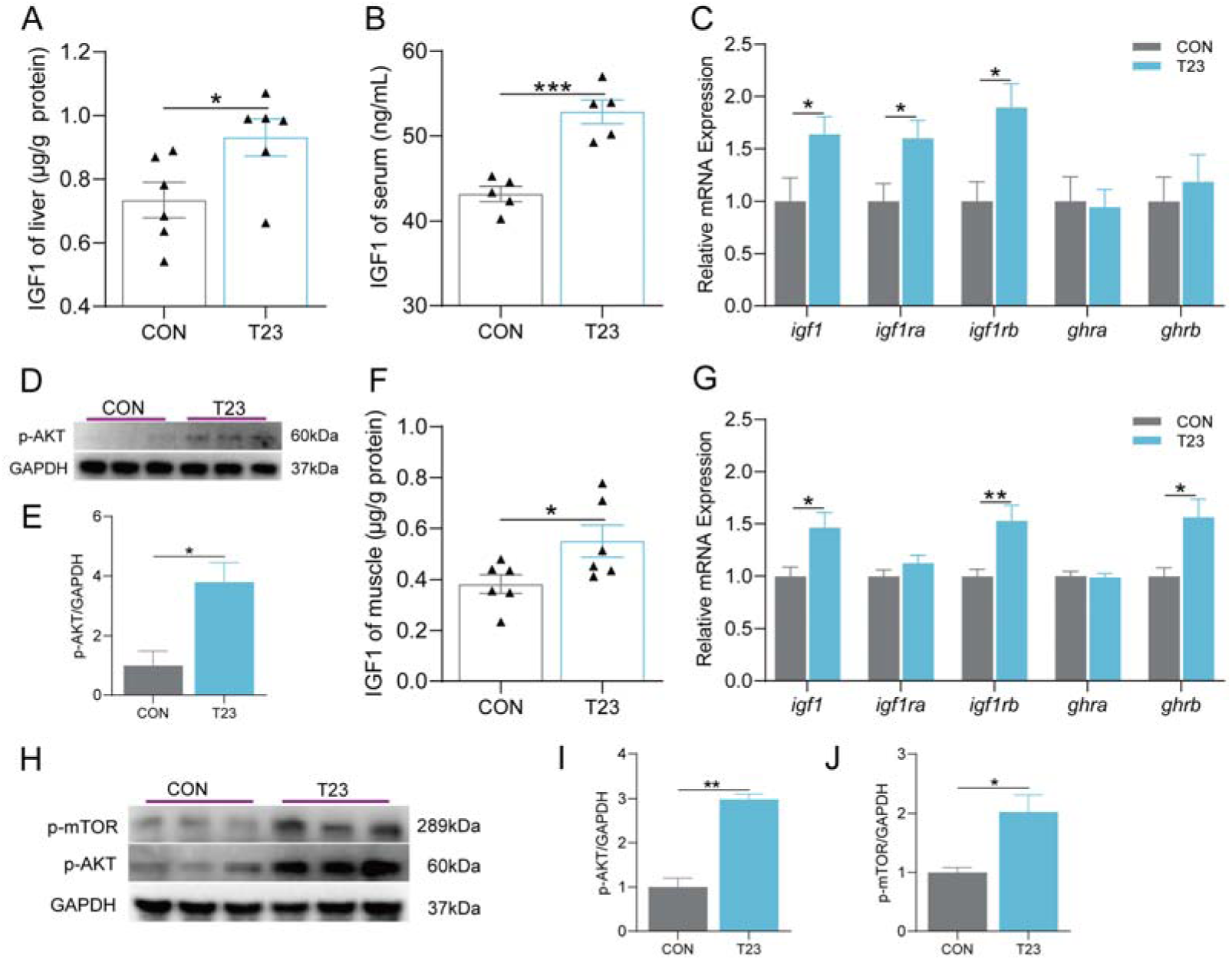
*B. velezensis* T23 promoted IGF1 signaling pathway of zebrafish. (A) The content of IGF1 in the liver (n = 6). (B) The content of IGF1 in the serum (n = 5). (C) The expression of growth-related genes in the liver (n = 6). (D) Western blot analysis and (E) quantitation of p-AKT in the liver (n = 3). (F) The content of IGF1 in the muscle (n = 6). (G) The expression of growth-related genes in the muscle (n = 6). (H) Western blot analysis of p-AKT and p-mTOR in the muscle (n = 3). (H) Quantitation of p-AKT in the muscle (n = 3). (H) Quantitation of p-mTOR in the muscle (n = 3). Data were presented as mean ± SEM. Student’s t-test was employed to compare between two groups. Significant differences were indicated as \**p* < 0.05, \*\**p* < 0.01, \*\*\**p* < 0.001.

### *B. velezensis* T23 modulated the gut microbiota of zebrafish

To evaluate the effect of *B. velezensis* T23 on the gut microbiota, 16S rRNA gene sequencing analysis of the gut content was conducted, and the results showed that the diversity and overall structure of the gut microbiota in the T23 treatment group did not significantly change versus CON group (Fig. 3A and B). However, in terms of the gut microbiota composition, T23 treatment led to changes in the relative abundance of some phyla and genera. The relative abundance of the Firmicutes was increased, while the relative abundances of Proteobacteria and its subordinate genera *Plesiomonas* and *Aeromonas* were decreased (Fig. 3C and D). Linear discriminant analysis effect size (LEfSe) further revealed that *B. velezensis* was a biomarker with significant difference in the T23 group (Fig. 3E), and its relative abundance in the T23 group was significantly higher than that in the CON group (Fig. 3F).

**Figure 3.**
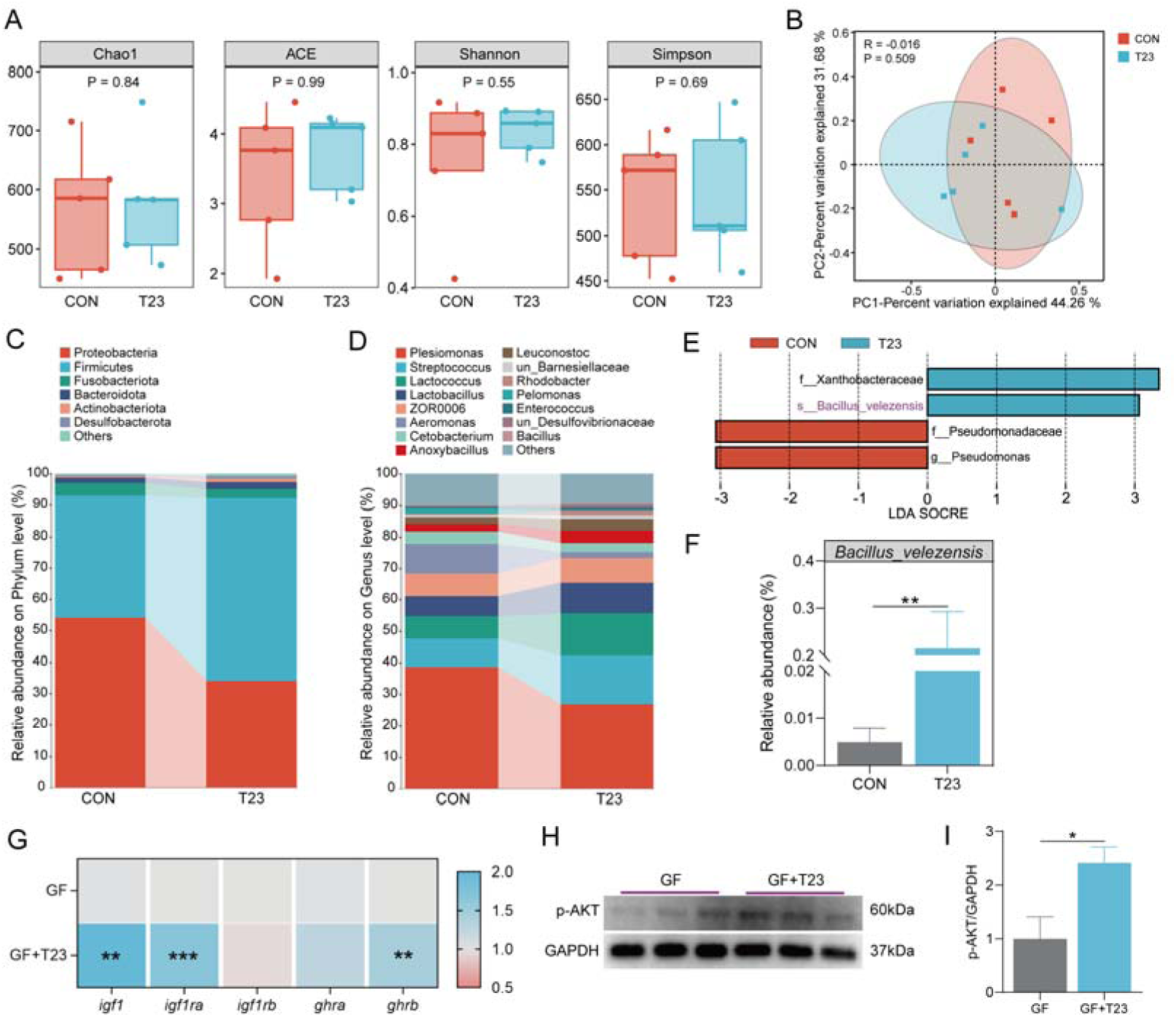
*B. velezensis* T23 directly activated the IGF1 signaling pathway in zebrafish. (A) Alpha diversity analysis of gut microbiota (n = 5). (B) PCoA analysis (Bray-Curtis) of gut microbiota at the OTU level (n = 5). (C) Relative abundance at the phylum level (n = 5). (D) Relative abundance at the genus level (n = 5). (E) LEfSe analysis of gut microbiota at the species level (n = 5). (F) The relative abundance of *B. velezensis* by using Mann-Whitney U test (n = 5). (G-I) GF zebrafish were divided into two groups: GF group (GF), and GF zebrafish mono-colonized with *B. velezensis* T23 (T23). Gnotobiotic zebrafish were fed with sterile diet for 7 days. (G) The expression of growth-related genes in gnotobiotic zebrafish (n = 4). (H) Western blot analysis and (I) quantitation of p-AKT in gnotobiotic zebrafish (n = 3). Data were presented as mean ± SEM. Student’s t-test was employed to compare between two groups. Significant differences were indicated as \**p* < 0.05, \*\**p* < 0.01, \*\*\**p* < 0.001.

### *B. velezensis* T23 directly promoted IGF1 production in gnotobiotic zebrafish

To investigate the microbiota-associated effect, the gut microbiota from CON or T23-fed zebrafish were transferred to germ-free (GF) zebrafish (Fig. S3A). The results showed that GF zebrafish colonized with T23 group-derived microbiota had significantly higher expression levels of *igf1*, *igf1ra*, *igf1rb*, *ghrb* compared with the counterparts colonized with the CON microbiota (Fig. S3B). As *B. velezensis* T23 colonized the gut after feeding, the effect of T23 group-derived microbiota might be due to the action of T23 *per se*. To prove this hypothesis, GF zebrafish was mono-colonized with *B. velezensis* T23, and results showed that the expression of *igf1*, *igf1ra*, *ghrb* and the phosphorylation of AKT in T23-colonized gnotobiotic zebrafish were significantly higher than that in GF zebrafish (Fig. 3G-I), indicating that *B. velezensis* T23 can directly interact with host to promote IGF1 production. To determine whether the effect on IGF1 in gnotobiotic zebrafish was specific to *Bacillus*, we mono-colonized GF zebrafish with other two commensal bacterial strains (*Plesiomonas shigeloides* CB5 or *Aeromonas veronii* XMX-5). Neither strain promoted IGF1 production in gnotobiotic zebrafish (Fig. S2C-G). Collectively, these results suggest that *B. velezensis* T23 can directly promote IGF1 signaling in zebrafish independent of the its modulatory role of the microbiota.

### The growth promoting effect of *B. velezensis* T23 was independent of lipoproteins and polyketides

*B. velezensis* produces abundant secondary metabolites (such as lipopeptides and polyketides), which are responsible for various beneficial functions [23]. To determine whether the growth-promoting effect of T23 depended on lipopeptides and polyketides, we compared the growth-promoting effects of WT *B. velezensis* T23 and T23-Δ*sfp*, a *sfp* gene knockout mutant strain deficient in the production of all lipopeptides and polyketides. Feeding experiment showed comparable improvement of FBW, WG, and FCR in zebrafish fed the T23 and T23-Δ*sfp* supplemented diet (Fig. 4A-C). Consistently, both T23 and T23-Δ*sfp* significantly increased the content of IGF1 in the liver and serum of zebrafish (Fig. 4D and E). Compared with the CON group, the expression of *igf1*, *igf1ra*, *igf1rb* in the liver of zebrafish was similarly increased after feeding with T23 and T23-Δ*sfp* (Fig. 4F), and the phosphorylation of AKT was significantly enhanced (Fig. 4G and H). Moreover, both T23 and T23-Δ*sfp* significantly increased the levels of IGF1 in the muscle of zebrafish (Fig. 4I). Compared with the control group, the expression of *igf1*, *igf1rb* in the muscle of zebrafish was increased after feeding with both T23 and T23-Δ*sfp* (Fig. 4J), and the phosphorylation of AKT and mTOR was similarly enhanced (Fig. 4K-L). Together, these results indicated that the growth promoting effect of T23 was not mediated by secondary metabolites lipopeptides and polyketides.

**Figure 4.**
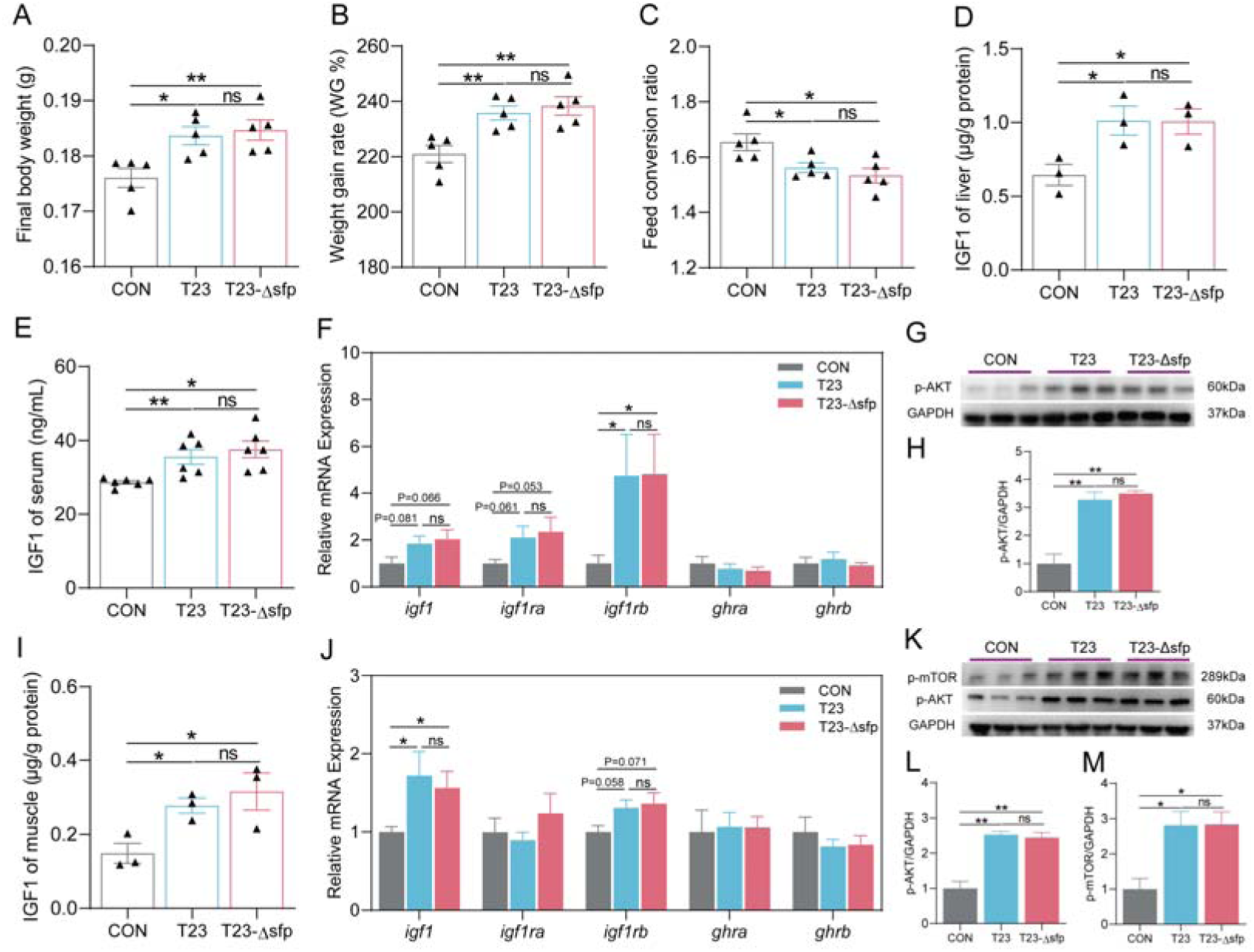
The growth promoting effect of *B. velezensis* T23 was independent of secondary metabolites lipopeptides and polyketides. One-month-old zebrafish were fed with normal diet (CON group) or diet supplemented with 10L CFUs/g *B. velezensis* T23 (T23 group) or *B. velezensis* T23-Δ*sfp* (T23-Δ*sfp* group) for 4 weeks. (A) Final body weight (FBW, n = 5). (B) Weight gain rate (WG, n = 5). (C) Feed conversion ratio (FCR, n = 5). (D) The content of IGF1 in the liver (n = 3). (E) The content of IGF1 in the serum (n = 5). (F) The expression of growth-related genes in the liver (n = 5). (G) Western blot analysis and (H) quantitation of p-AKT in the liver (n = 3). (I) The content of IGF1 in the muscle (n = 3). (J) The expression of growth-related genes in the muscle (n = 5). (K) Western blot analysis of p-AKT and p-mTOR in the muscle (n = 3). (L) Quantitation of p-AKT in the muscle (n = 3). (M) Quantitation of p-mTOR in the muscle (n = 3). Data were presented as mean ± SEM. Student’s t-test was employed to compare between two groups. Significant differences were indicated as \**p* < 0.05, \*\**p* < 0.01.

### *B. velezensis* T23 isolated cell wall peptidoglycan was sufficient to promote growth and IGF1 production

To explore whether the cell wall components of *B. velezensis* T23 mediated the growth-promoting effect, we extracted the whole cell wall and purified peptidoglycan of *B. velezensis* T23, and conducted a 4-week feeding experiment to evaluate the growth-promoting effect of the whole cell wall (WCW) and peptidoglycan (PGN). During the experiment, there was no significant difference in the survival rate of zebrafish groups (Fig. S3A). Compared with the CON group, dietary supplementation of cell wall or peptidoglycan significantly increased the FBW and WG of zebrafish (Fig. S3B and 5A), and significantly reduced the FCR (Fig. 5B). In terms of physical indicators, there was no significant difference in the body length of zebrafish between the two treatment groups compared to the CON group (Fig. S3C), but the CF was significantly increased (Fig. S3D). The indicators of edible part showed that compared with the CON group, the VSI in the WCW and PGN groups tended to decrease (Fig. S3E), and the CR tended to increase (Fig. S3F). Muscle component analysis showed that the muscle moisture of the WCW and PGN groups was significantly reduced (Fig. S3G), and the protein content was significantly increased (Fig. S3H). Consistent with live bacterium, both the whole cell wall and peptidoglycan significantly increased the IGF1 levels in the liver and serum of zebrafish (Fig. 5C and D), and significantly up-regulated the gene expression of *igf1*, *igf1ra*, *igf1rb* in the liver (Fig. 5E). In muscle, both components significantly increased the IGF1 levels (Fig. 5F), up-regulated the gene expression of *igf1*, *igf1ra* (Fig. 5G), and significantly enhanced the phosphorylation of AKT and mTOR (Fig. 5H-J). Therefore, the cell wall and peptidoglycan were sufficient to promote growth and IGF1 production in zebrafish, suggesting that the growth-promoting effect of *B. velezensis* T23 was mediated by cell wall component, and more specifically the peptidoglycan.

**Figure 5.**
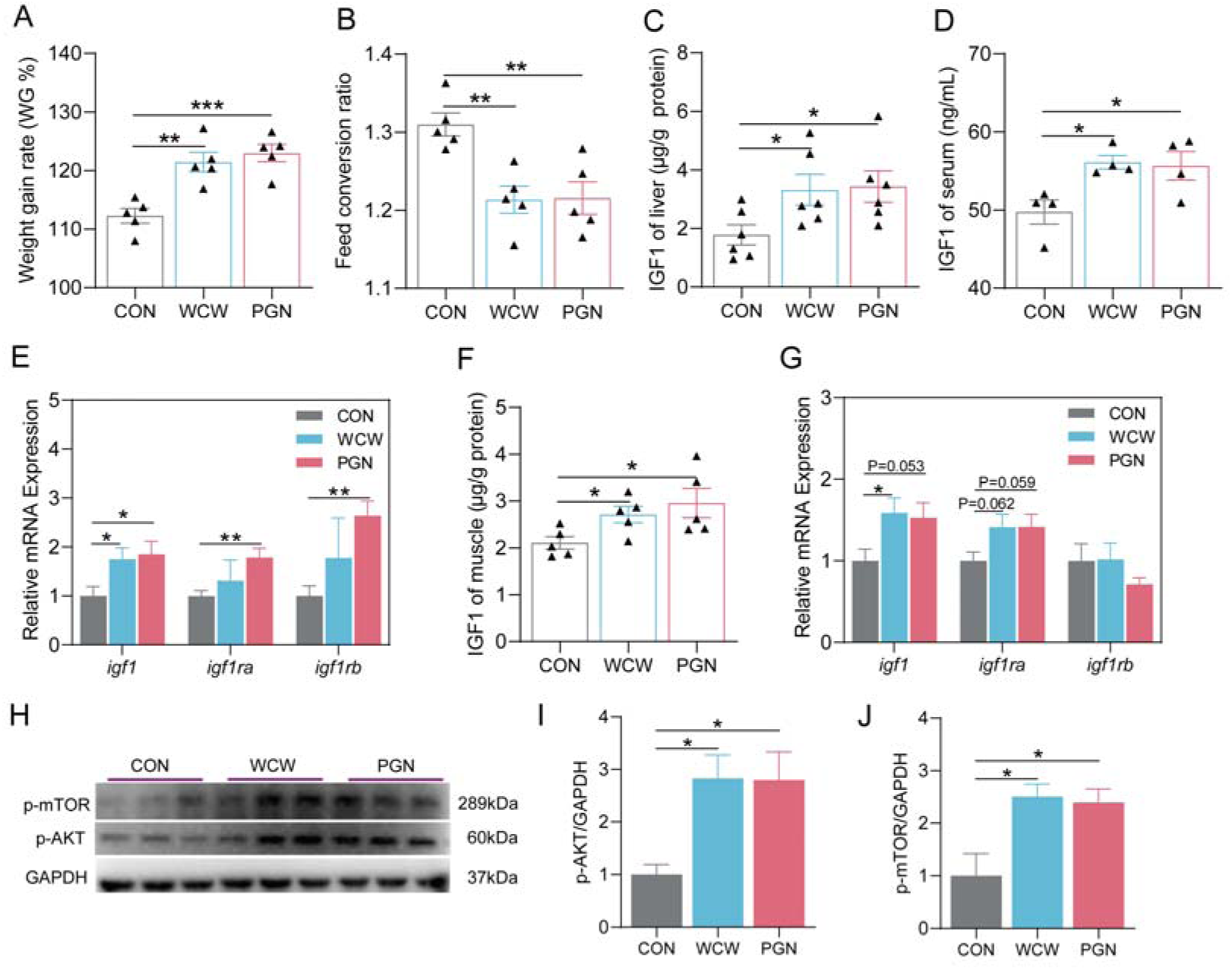
Cell wall peptidoglycan mediated the stimulatory effect of *B. velezensis* T23 on IGF1 signaling pathway. One-month-old zebrafish were respectively fed with normal diet (CON group) or diet supplemented with equal amounts of the whole cell wall (WCW group) or purified peptidoglycan (PGN group) of *B. velezensis* T23 (10L CFUs/g) for 4 weeks. (A) Weight gain rate (WG, n = 5). (B) Feed conversion ratio (FCR, n = 5). (C) The content of IGF1 in the liver (n = 6). (D) The content of IGF1 in the serum (n = 4). (E) The expression of growth-related genes in the liver (n = 5). (F) The content of IGF1 in the muscle (n = 5). (G) The expression of growth-related genes in the muscle (n = 5). (H) Western blot analysis of p-AKT and p-mTOR in the muscle (n = 3). (I) Quantitation of p-AKT in the muscle (n = 3). (J) Quantitation of p-mTOR in the muscle (n = 3). Data were presented as mean ± SEM. Student’s t-test was employed to compare between two groups. Significant differences were indicated as \**p* < 0.05, \*\**p* < 0.01, \*\*\**p* < 0.001.

### Cell wall-derived muramyl dipeptide (MDP) promoted IGF1 production and growth in zebrafish

Peptidoglycan hydrolysis generates PGN fragments known as muropeptides. As the minimal functional component of PGN, MDP exhibited various beneficial effects on the immunity and health of host [30, 31]. Therefore, we further investigated the effect of MDP on growth and IGF1 signaling in zebrafish. Firstly, we observed that the content of MDP in the intestinal contents of zebrafish in the group fed with *B. velezensis* T23 was significantly higher than that of the CON group (Fig. 6A). Under *in vitro* conditions, *B. velezensis* T23 was able to produce MDP (Fig. S4A). Additionally, the number of *B. velezensis* was significantly positively correlated with MDP content in the intestinal contents (Fig. S4B). Furthermore, dietary inclusion of the optimal dose of MDP significantly increased FBW and WG of zebrafish (Fig. S5A and 6B), and reduced FCR (Fig. 6C). No significant difference was observed in body length between the two groups (Fig. S5B), but the CF was significantly increased (Fig. S5C). The edible-part indicators showed that compared with the CON group, the VSI of zebrafish in the MDP group was decreased (Fig. S5D), and the CR was significantly increased (Fig. S5E). The muscle moisture of MDP group was significantly reduced (Fig. S5F), and the protein content was significantly increased (Fig. S5G).

**Figure 6.**
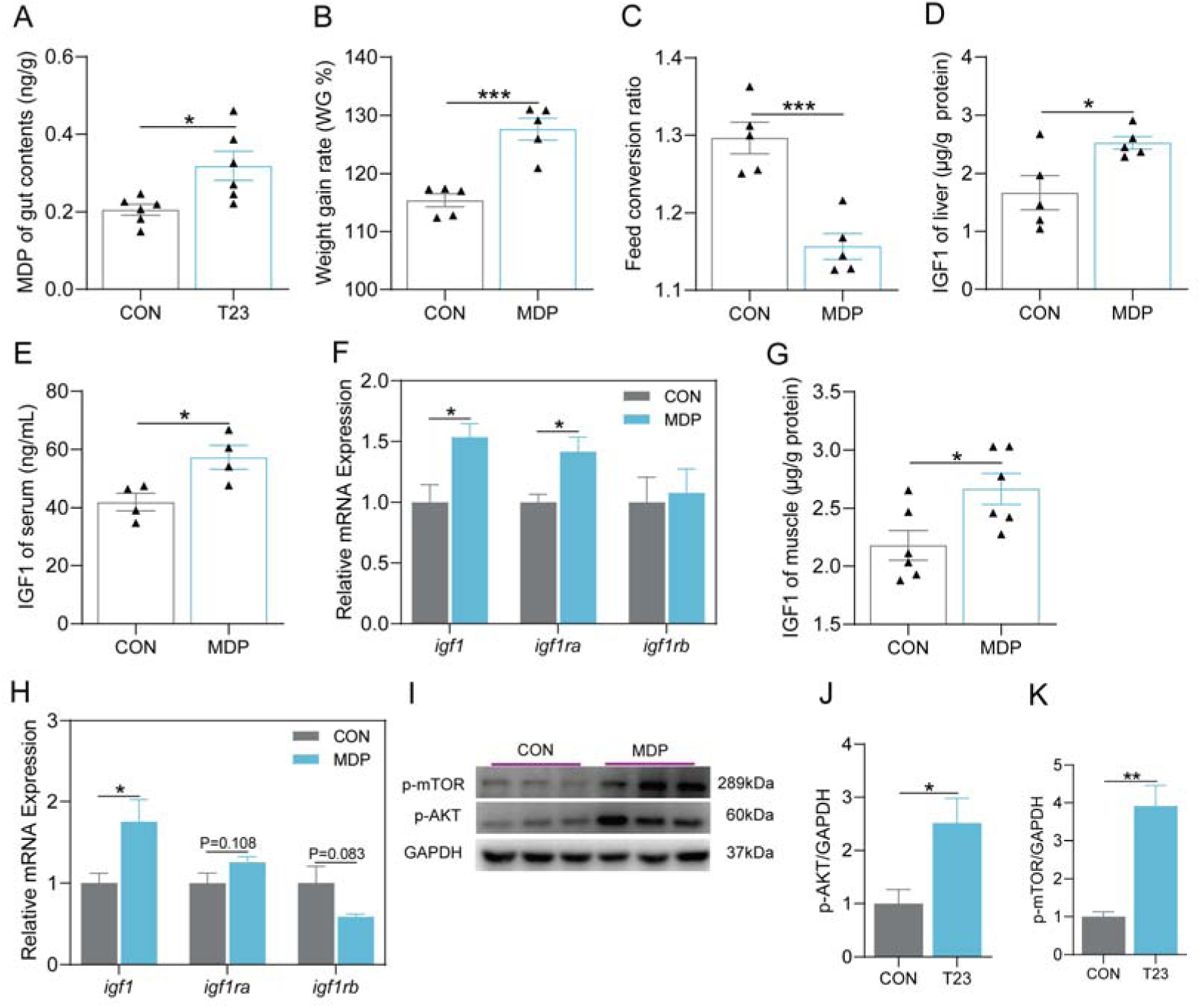
MDP was sufficient to activate the IGF1 signaling pathway in zebrafish. (A) The content of MDP in the intestinal contents of zebrafish (n = 6). One-month-old zebrafish were fed with normal diet (CON group) or diet supplemented with 0.00015% MDP (MDP group) for 4 weeks. (B) Weight gain rate (WG, n = 5). (C) Feed conversion ratio (FCR, n = 5). (D) The content of IGF1 in the liver (n = 5). (E) The content of IGF1 in the serum (n = 4). (F) The expression of growth-related genes in the liver (n = 5). (G) The content of IGF1 in the muscle (n = 6). (H) The expression of growth-related genes in the muscle (n = 5). (I) Western blot analysis of p-AKT and p-mTOR in the muscle (n = 3). (J) Quantitation of p-AKT in the muscle (n = 3). (K) Quantitation of p-mTOR in the muscle (n = 3). Data were presented as mean ± SEM. Student’s t-test was employed to compare between two groups. Significant differences were indicated as \**p* < 0.05, \*\**p* < 0.01, \*\*\**p* < 0.001.

Consistent with the effects of *B. velezensis* T23 and peptidoglycan, MDP significantly increased IGF1 levels in the liver and serum of zebrafish (Fig. 6D and E). Compared with the CON group, the gene expression of *igf1*, *igf1ra* in the liver of zebrafish was increased significantly after feeding MDP (Fig. 6F). In addition, MDP could significantly increase IGF1 level (Fig. 6G) and the gene expression of *igf1*, *igf1ra* in the muscle of zebrafish (Fig. 6H), with increased AKT and mTOR phosphorylation (Fig. 6I-K). Together, the results indicated that cell wall-derived MDP recapitulated the effect and action mode of *B. velezensis* T23 on growth and IGF1 production in zebrafish, supporting that the growth promoting effect of T23 was mediated by cell wall derived MDP.

### *B. velezensis* T23 promoted IGF1 production and growth of zebrafish via NOD2 signaling

NOD2 is the canonical receptor of MDP [8, 30]. Therefore, we further verified the necessity of NOD2 in the growth promoting effect of *B. velezensis* T23. Firstly, in zebrafish larvae treated with vivo morpholino directed to NOD2, *B. velezensis* T23 no longer increased the IGF1 content and the expression of its receptor genes (Fig. S6A-D), and the increase in AKT phosphorylation was abrogated (Fig. S6C and D). Further, we used CRISPR/Cas9 to construct *nod2*^-/-^ zebrafish (Fig. 7A and B). The results showed that *B. velezensis* T23 treatment had no significant effect on the WG and FCR of *nod2*^-/-^ zebrafish (Fig. 7C and D), and the effect of *B. velezensis* T23 on IGF1 production in the liver and muscle was blocked (Fig. 7E and F). Similarly, the phosphorylation of AKT and mTOR was not activated by *B. velezensis* T23 in *nod2*^-/-^ zebrafish (Fig. 7G-I). No significant difference in the colonization of *B. velezensis* and MDP content in the intestine of WT and *nod2*^-/-^ zebrafish (Fig. 7J and K), excluding that the negative phenotypes observed in *nod2*^-/-^zebrafish was due to differential T23 colonization or MDP production. Together, these results confirmed that MDP-NOD2 signaling was necessary for the effect of *B. velezensis* T23 on growth and IGF1 production in zebrafish.

**Figure 7.**
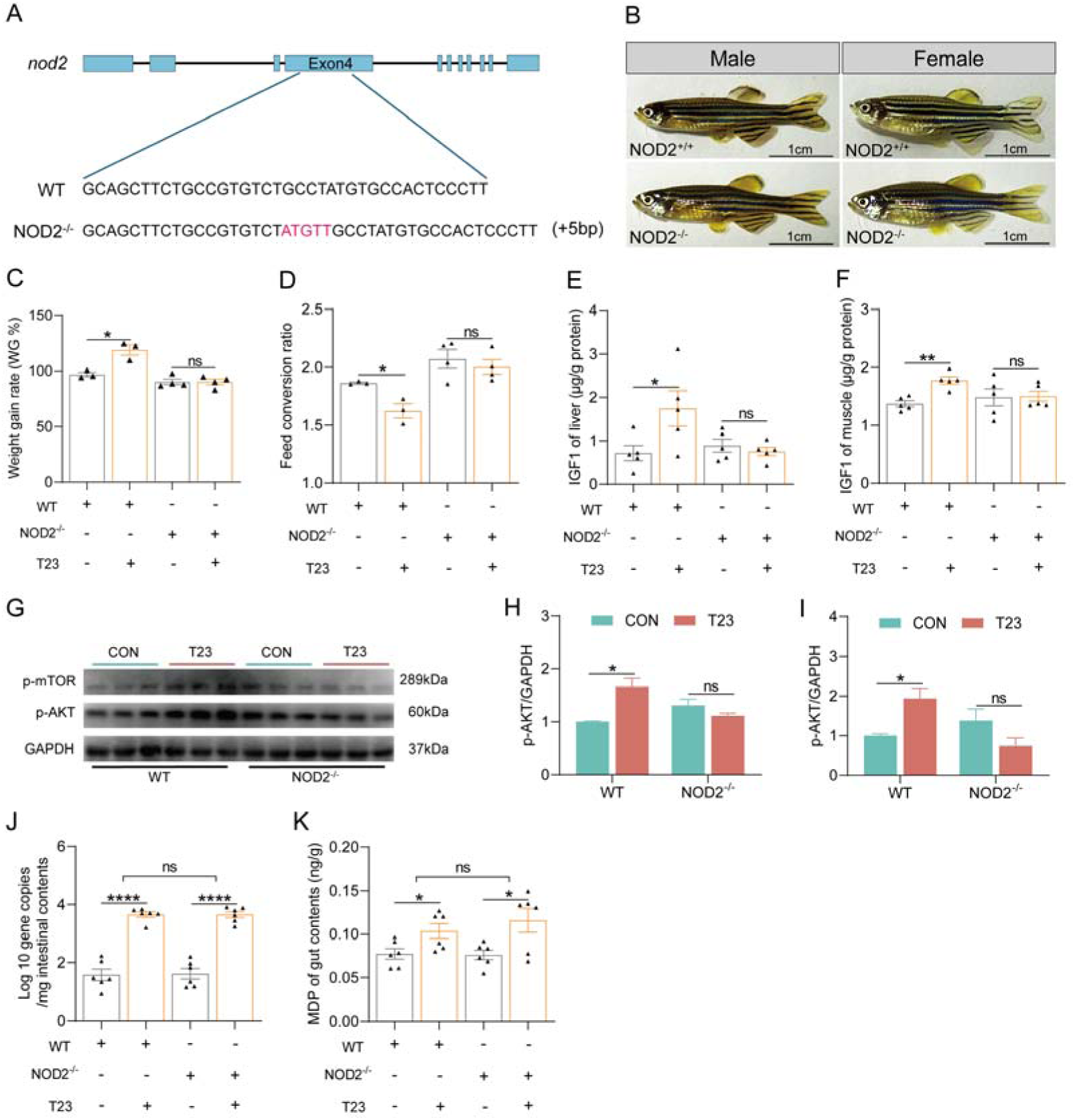
*B. velezensis* T23 stimulated IGF1 production via NOD2 signaling. (A) Schematic diagram of CRISPR/Cas9 editing targeting exon 4 of the *nod2* gene. (B) Comparison of the appearance between WT and *nod2*^-/-^ zebrafish (male; female). (C-K) One-month-old zebrafish (WT and *nod2*^-/-^) were fed with normal diet or diet supplemented with 10L CFUs/g *B. velezensis* T23 for 4 weeks. (C) Weight gain rate (WG, n = 3-4). (D) Feed conversion ratio (FCR, n = 3-4). (E) The content of IGF1 in the liver (n = 5). (F) The content of IGF1 in the muscle (n = 5). (G) Western blot analysis of p-AKT and p-mTOR in the muscle (n = 3). (H) Quantitation of p-AKT in the muscle (n = 3). (I) Quantitation of p-mTOR in the muscle (n = 3). (J) The copies of *B. velezensis* in the intestinal contents (n = 6). (K) The content of MDP in the intestinal contents of zebrafish (n = 6). Data were presented as mean ± SEM. Student’s t-test was employed to compare between two groups. Significant differences were indicated as \**p* < 0.05, \*\**p* < 0.01, \*\*\*\**p* < 0.0001, ns=not significant.

### *B. velezensis* T23 promoted the renewal and differentiation of intestinal cells

Through *in vitro* experiments in ZFL cells, it was found that MDP was not able to directly increase the production of IGF1 in ZFL cells and the expression of related receptor genes (Fig. S7A-D). This indicated that MDP did not directly act on the liver and the action was probably mediated via gut-liver axis. To further explore how host intestinal cells perceive and respond to MDP, we conducted transcriptome sequencing analysis of zebrafish intestinal tissues to investigate pathways downstream of NOD2 signaling. The overall gene expression levels of each sample were basically the same (Fig. S8A). Principal component analysis showed that the samples in the T23 group were significantly separated from the CON group (Fig. 8A), indicating that T23 significantly changed the gene expression profile of the intestine. A total of 666 differentially expressed genes were identified between the two groups, with 363 upregulated and 303 downregulated, and the two most significant genes were *hif1*α and *notch3* (Fig. 8B and S8B). Functional enrichment analysis revealed the extensive impact of T23 treatment on intestinal physiological processes. KEGG pathway enrichment analysis showed that T23 treatment significantly affected multiple signaling pathways closely related to cellular processes (Fig. S8C), among which the cell cycle pathway was the most significantly upregulated (Fig. 8C). GO enrichment analysis indicated that T23 significantly changed the number of genes related to biological process and cellular component (Fig. S8D). Gene set enrichment analysis (GSEA) further confirmed that T23 treatment significantly activated related pathways such as cell cycle (Fig. 8D), DNA replication (Fig. 8E), and homologous recombination (Fig. 8F). Combined with weighted gene co-expression network analysis (WGCNA), based on the gene expression characteristics of individual samples, clustering was performed (Fig. 8G), and the gene expression in the brown module was distinct from that of the CON group and was upregulated in the intestinal tissues of the T23 group (Fig. 8H). Further KEGG functional enrichment analysis of the genes within the brown module showed that the most significantly upregulated signaling pathway was also cell cycle (Fig. 8I). These results suggest that T23 treatment may promote the renewal and differentiation of intestinal cells. Histological analysis further confirmed this speculation: compared with the CON group, the height of intestinal villus was significantly increased in the T23 group (Fig. S9A and B), and the number of goblet cells was also significantly increased (Fig. S9C).

**Figure 8.**
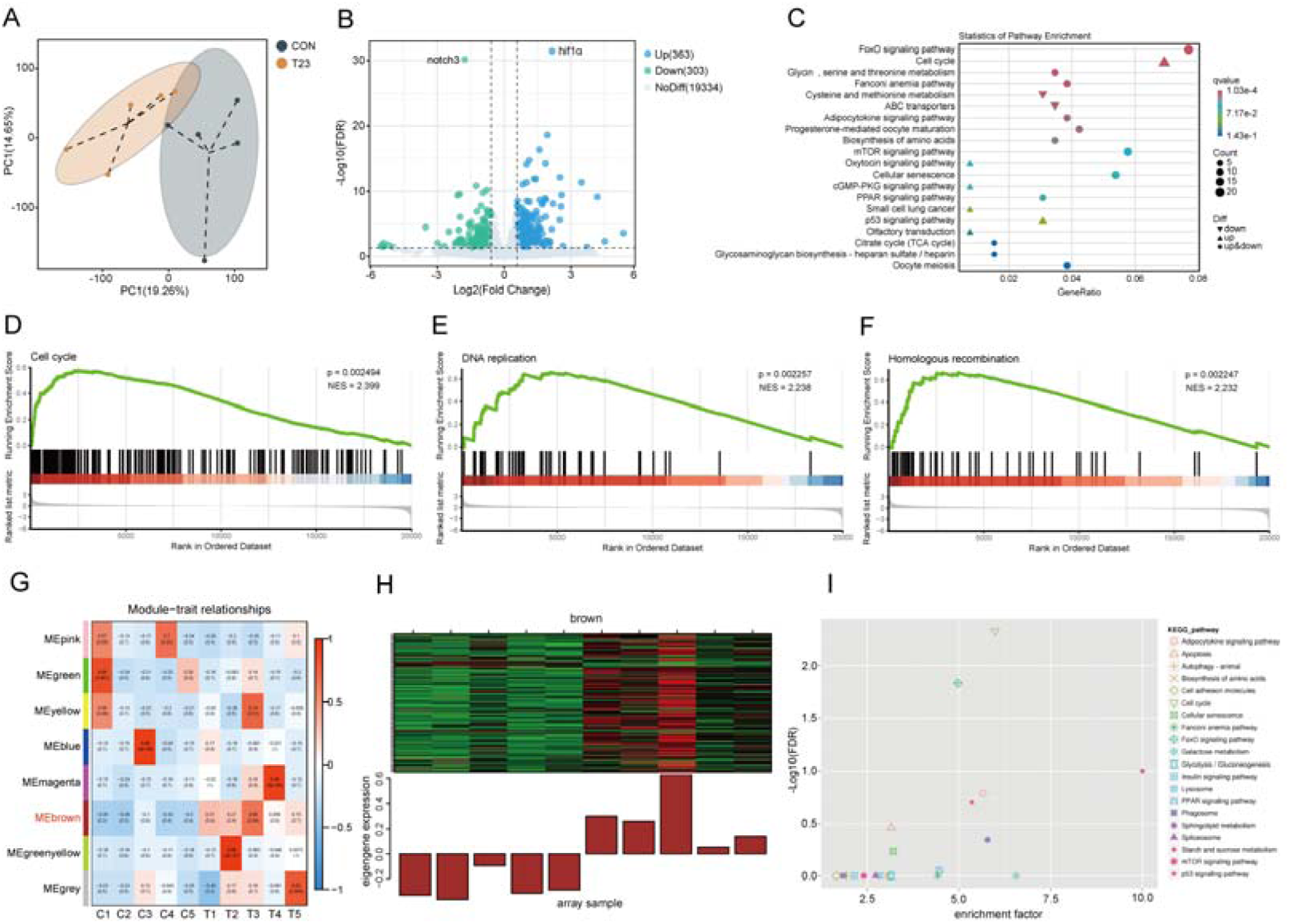
*B. velezensis* T23 promoted the renewal and differentiation of intestinal cells. (A) PCA diagram of various transcriptome samples (n = 5). (B) The volcano plot showed the overall distribution of differentially expressed genes (DEGs) (Fold Change ≥ 1.5, FDR < 0.05). (C) KEGG enrichment analysis of DEGs. (D) GSEA of the cell cycle pathway. (E) GSEA of the DNA replication pathway. (F) GSEA of the homologous recombination pathway. (G) The co-expression module of WGCNA. (H) Gene expression in the brown module of WGCNA. (I) KEGG enrichment analysis in the brown module of WGCNA.

By integrating the results of these multi-dimensional analyses, we focused on the cell cycle pathway and its upstream regulatory factors. Compared with the CON group, the expression of *hif1*α and cell replication-related genes (*ccnb1*, *ccnb2*, *ccnb3*, *cdk1*, *ccne2*, *ttk*, *plk1*, *skp2*, *anapc1*, *gadd45ab*, *gadd45ba* and *gadd45ga*) was significantly upregulated in the T23 group, while the expression of genes (*notch3*, *her6* and *hey1*) related to the inhibition of cell differentiation was significantly decreased (Fig. S8E). Previous study showed that HIF1a can activate the cell cycle pathway [32]. Therefore, T23 might activate the cell replication-related pathways of intestinal cells via HIF1a signaling, and promote intestinal cell renewal and improvement of the intestinal villus structure, which may improve nutrient absorption, leading to increased activity of GH/IGF1 axis.

## Discussion

In this study, we found that the growth promoting effect of *B. velezensis* T23 was mediated by peptidoglycan-derived MDP, which engaged NOD2 to stimulate IGF1 production. The overall impact of the microbiota on host physiology has been reported in previous studies. Yan et al. demonstrated that the colonization of a normal microbiota could induce the production of IGF1 in conventional mice and promote bone formation, while germ-free (GF) mice showed decreased circulating IGF1 levels and growth retardation [33], confirming the importance of the gut microbiota in promoting host growth. Similarly, the growth performance of conventional mice was significantly higher than that of GF mice [5]. The bacterial taxa with the stimulatory effect on IGF1 production have also been studied. Commensal *P. copri* induced the production of IL-13 in the intestine, which entered the liver via the portal vein and activates the IL-13R/Jak2/Stat6 pathway, thereby promoting the synthesis of IGF1 in the liver of mice and pigs [34]. Human breastmilk-derived *Bifidobacterium longum* regulated the composition of the gut microbiota and its metabolites to activate the GH/IGF-1 axis and the downstream PI3K/AKT signaling pathway, thereby improving bone metabolism during the growth process [35]. In a recent pioneered study, Schwarzer et al. reported that the cell walls of *L. plantarum* were able to sustain the growth of chronically malnourished mice and supports the IGF1 production, which was mediated by cell wall-derived MDP mediated NOD2 stimulation [8]. Our results showed that PG-derived MDP from another species of Firmicutes can stimulate growth via NOD2-IGF1 signaling in normally feeding zebrafish, suggesting that commensal bacterial PG-derived MDP has a conserved effect on IGF1 production in vertebrates irrespective of nutritional status.

The effector molecules of the gut microbiota can directly act on distal organs or mediate signal transmission through receptors on intestinal cells [24, 26, 36, 37]. In this study, *in vitro* experiments using ZFL cells model showed that MDP was not able to directly increase the IGF1 content and the expression of receptor genes in ZFL cells (Fig. S7A-D), indicating that MDP did not directly act on the NOD2 on liver cells. Consistently, the growth promoting effect of *L. plantarum* depended on intestinal NOD2 signaling, but not NOD2 in hepatocytes [8], indicating that PG-derived MDP engaged NOD2 in the intestinal epithelial cells to stimulate IGF1 production in the liver. One remaining question is how intestinal NOD2 stimulation induces liver IGF1 production. Schwarzer et al. proposed that *L. plantarum*-mediated NOD2 signaling in the crypt improved intestinal cell proliferation, epithelium maturation and thus nutrient absorption, thereby increasing the activity of the nutrient-sensitive GH/IGF1 axis [8]. In line with this, Nigro et al. reported that MDP activated NOD2 in Lgr5(+) stem cells at the intestinal crypt, thereby promoting cell survival [38, 39]. In this study, we also observed that *B. velezensis* promoted intestinal cell renewal and differentiation (Fig. 8C-I), and the intestinal villus structure was improved. Intriguingly, *B. velezensis* upregulated *hif1a* and downregulated *notch3* expression in the intestine (Fig. 8B and S8E). Previous studies have shown that HIF1α regulates the expression of various intestinal hormones, enhances the intestinal barrier function, and participates in nutrient absorption and metabolism [40, 41]. Therefore, the downstream pathway that mediates NOD2-induced IGF1 production may involve HIF1α signaling, which deserves further investigation.

Commensal bacteria may also regulate IGF1 production in a NOD2-independent way. For instance, microbiota-derived *P. copri* induces the transformation of intestinal CD4^+^ T cells into T helper 2 cells, promoting the secretion of IL-13 into the portal circulation, and subsequently stimulating IGF-1 production by activating the IL-13R/Jak2/Stat6 pathway [34]. In the above study, T cells were involved in IGF1 stimulation mediated by commensal bacterium or metabolites. Notably, in our study, zebrafish larvae were able to replicate the phenotype of IGF1 upregulation, indicating that it was not dependent on adaptive immunity. The metabolites of the microbiota could directly enter the circulation to stimulate IGF1 production in the liver. Yang et al. found that *Lactiplantibacillus plantarum-*5-hydroxyindole-3-acetic acid activated the hepatic AhR, which then promoted the production of IGF1 through the IRS1/PI3K/AKT pathway [42]. Furthermore, previous study found that a commensal *Escherichia coli* strain sustained IGF1 production of mice in disease conditions via direct engagement of the NLRC4 inflammasome in white adipose tissue [43]. Although this effect did not involve an action of the commensal bacterium in gut and liver, it implied that other members of the NLR family may also be involved in the regulation of systemic IGF1 signaling by gut bacteria. It will be interesting to identify other commensal taxa with NOD2-independent IGF1 stimulatory mode, and to investigate the contribution of NOD2 signaling to the overall effect of gut microbiota on IGF1 by using NOD2 KO GF animals.

## Acknowledgements

This study was funded by National Natural Science Foundation of China (NSFC 32330110, 31602169).

## Author contributions

All authors approved the final manuscript. C.R. and Z. Zhou designed the experiments; D.M. conducted the experiment and analyzed the data; W.Z., H.L., S.X., Y.Z., Y.W., Y. Yang, Z. Zhang, Y. Yao., Q.D. M.L., N.W., C.W. and Y.T. helped with the experiments or provided the reagents; D.M. and C.R. prepared the manuscript.

## Data availability

Any additional information required to reanalyze the data reported in this article can be obtained from the corresponding authors upon reasonable request.

