## Supplementary Materials for "*Bacillus velezensis*-derived muropeptide promotes growth of zebrafish via NOD2-mediated induction of IGF1 signaling"

**Supplemental Figures**

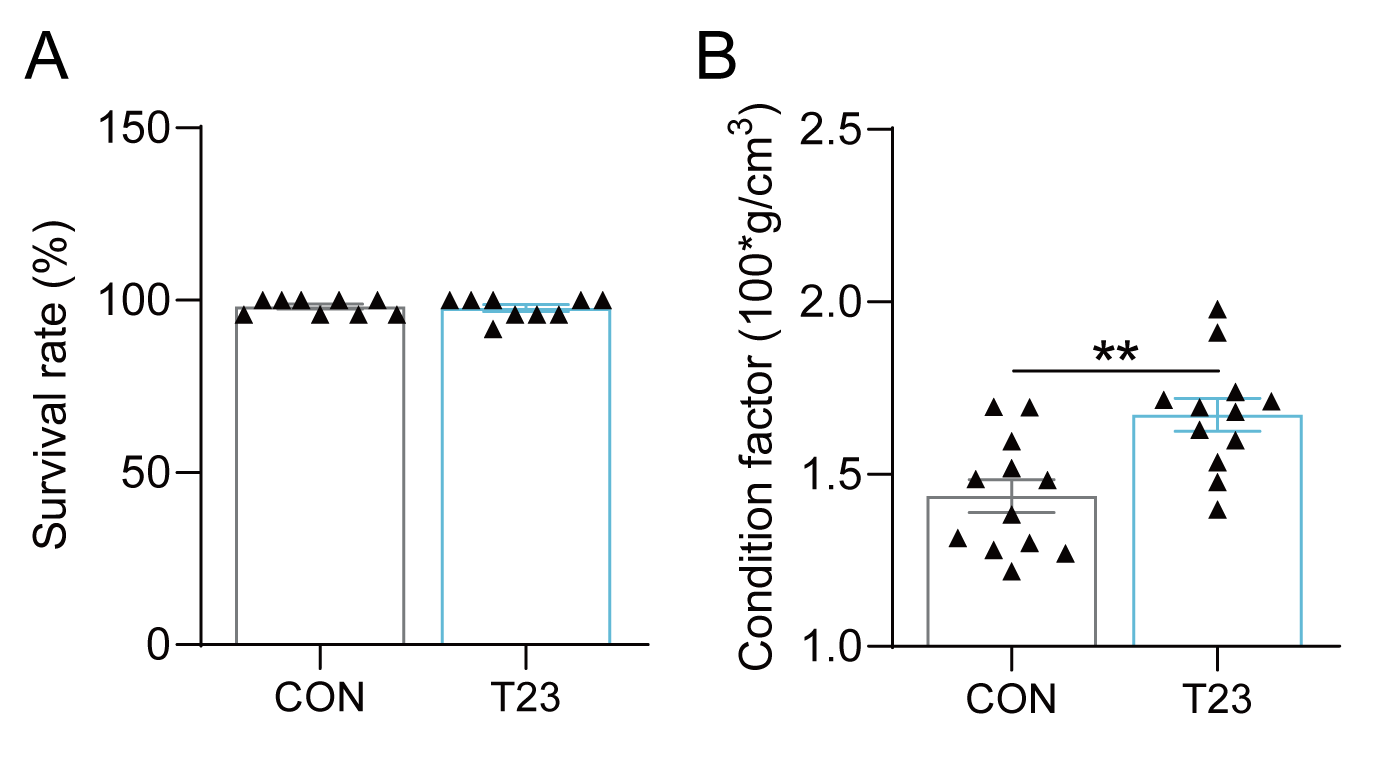

**Figure S1.** The effects of *B. velezensis* T23 on survival and CF of zebrafish. (A) Survival rate (SR, n = 9). (B) Condition factor (CF, n = 9). Data were presented as mean ± SEM. Student’s t-test was employed to compare between two groups. Significant differences were indicated as ***p* < 0.01.

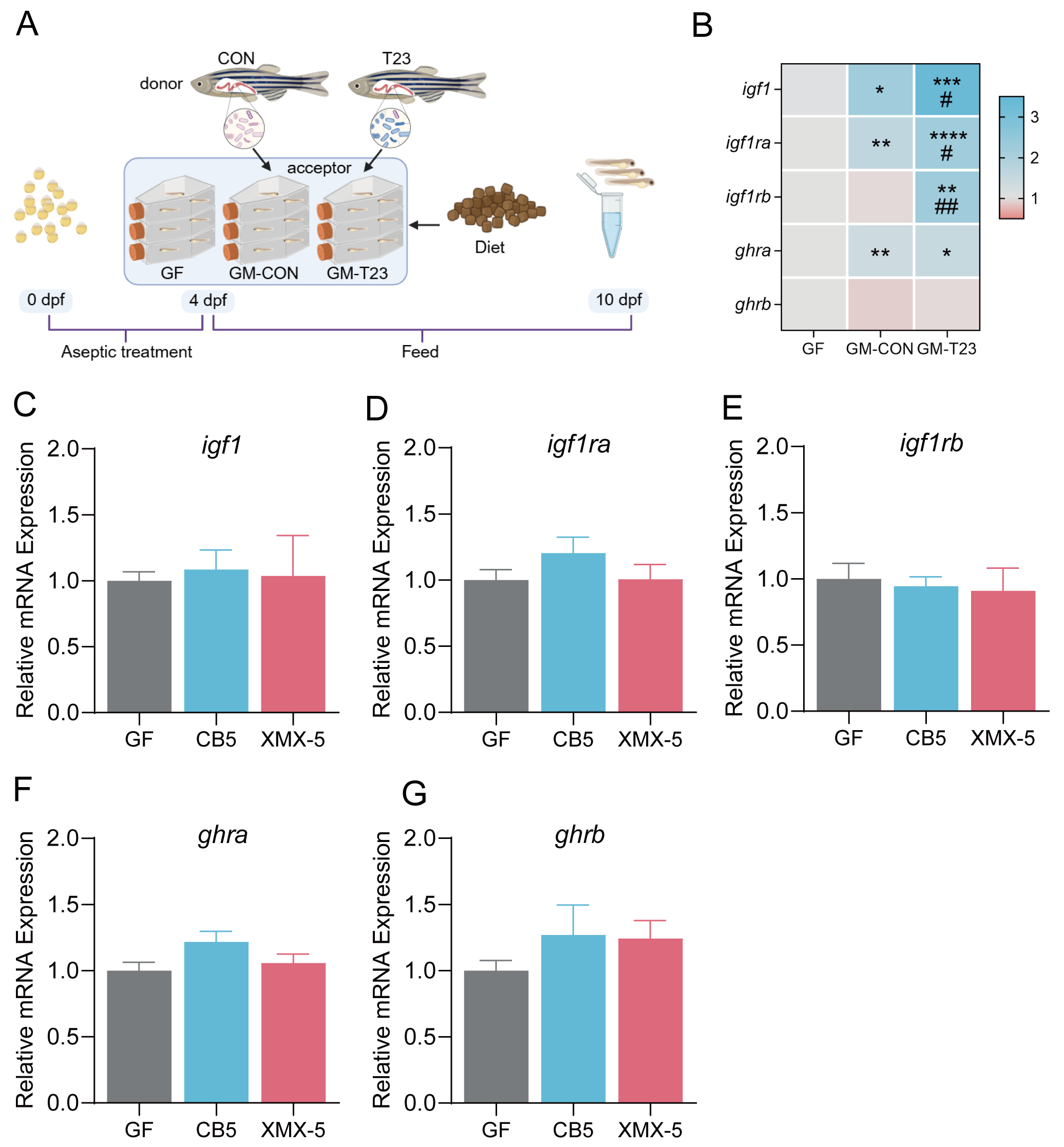

**Figure S2.** The effects of transferring gut microbiota or bacterium strain on the IGF1 signaling pathway in GF zebrafish. (A) The GF zebrafish larvae (6 replicate bottles per group, 30 fish per bottle) were divided into three groups: the GF zebrafish larvae group (GF), the GF zebrafish colonized with gut microbiota from CON group (GM-CON), or gut microbiota from T23 group (GM-T23). All the zebrafish larvae were fed with sterile diet for 7 days. (B) The expression of growth-related genes in zebrafish larvae (n = 6). The GF zebrafish larvae (4 replicate bottles per group, 30 fish per bottle) were divided into three groups: GF zebrafish (GF), and GF zebrafish mono-colonized with *P. shigelloides* CB5 (CB5) or *A. veronii* XMX-5 (XMX-5). All the zebrafish larvae were fed with sterile diet for 7 days. (C-G) The expression of growth-related genes in zebrafish larvae (n = 4). Data were presented as mean ± SEM. Student’s t-test was employed to compare between two groups. When the control group was GF, significant differences were indicated by **p* < 0.05, ***p* < 0.01, ****p* < 0.001, *****p* < 0.0001. When the control group was GM-CON, significant differences were indicated by ^#^*p* < 0.05, ^##^*p* < 0.01.

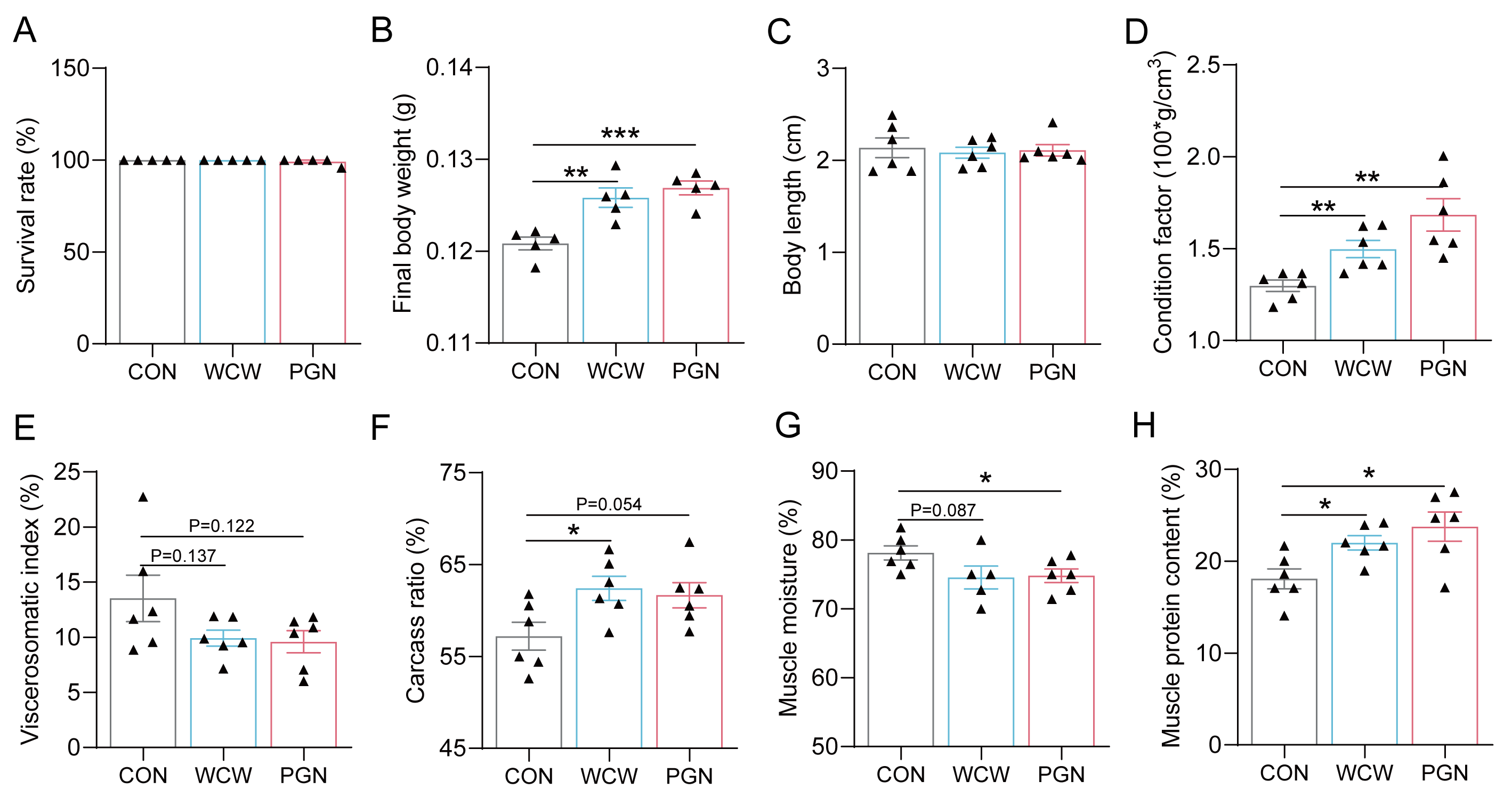

**Figure S3.** The effects of dietary supplementation of whole cell wall or peptidoglycan on the growth performance of zebrafish. (A) Survival rate (SR, n = 5). (B) Final body weight (FBW, n = 5). (C) Body length (n = 6). (D) Condition factor (CF, n = 6). (E) Viscerosomatic index (VSI, n = 6). (F) Carcass ratio (CR, n = 6). (G) Muscle moisture (n = 6). (H) Muscle protein content (n = 6). Data were presented as mean ± SEM. Student’s t-test was employed to compare between two groups. Significant differences were indicated as **p* < 0.05, ***p* < 0.01, ****p* < 0.001.

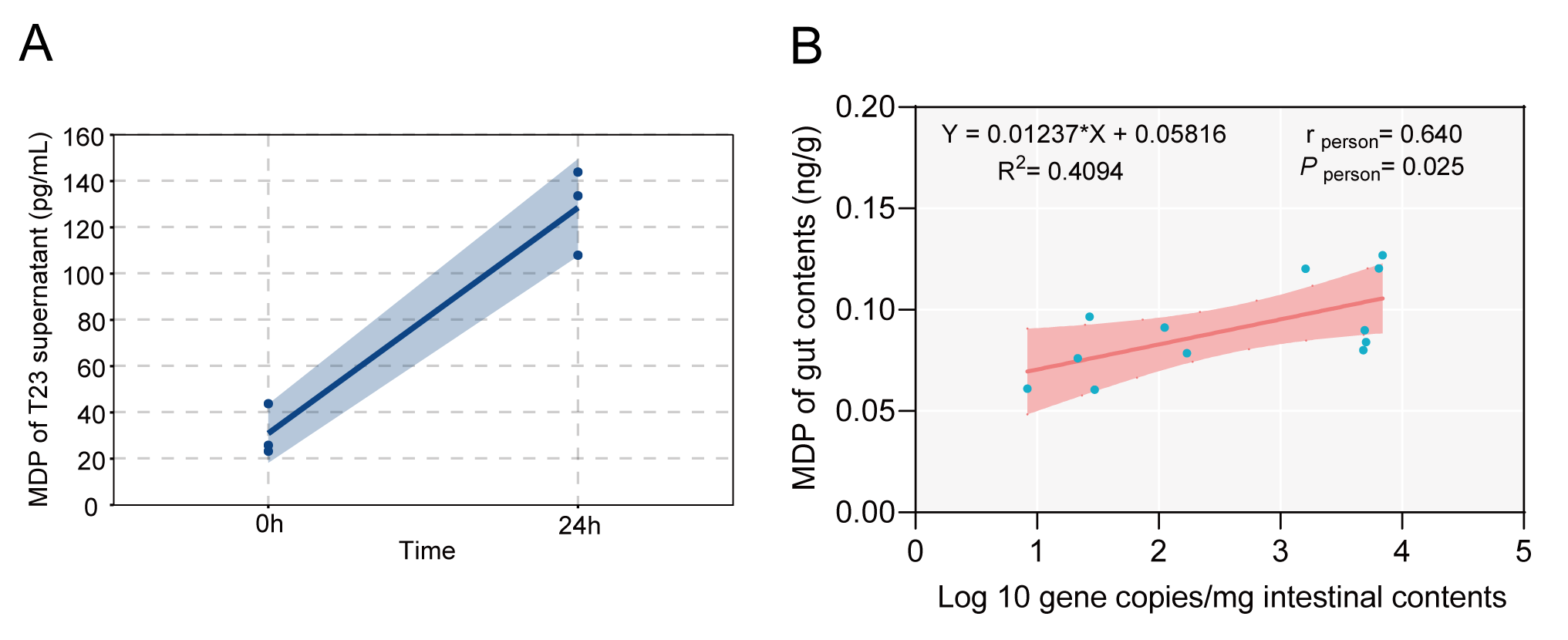
**Figure S4.** MDP content in *B. velezensis* culture or intestinal contents of zebrafish. (A) The MDP content in the supernatant of *B. velezensis* T23 (n = 3). (B) Pearson correlation between the copies of *B. velezensis* and the content of MDP in the intestinal contents of zebrafish (n = 12).

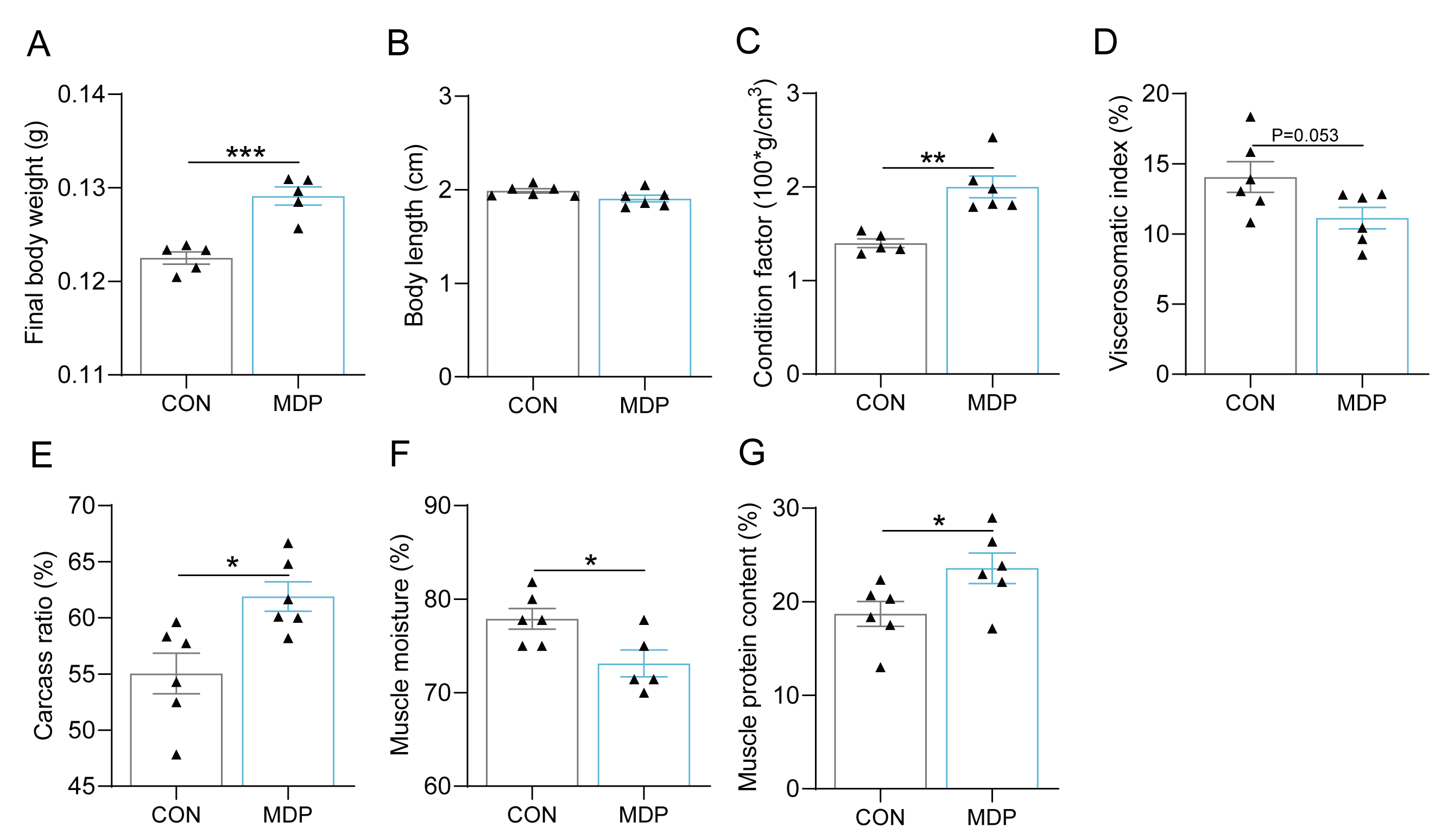

**Figure S5.** The effects of dietary supplementation of MDP on IGF1 signaling and growth of zebrafish larvae. Zebrafish larvae were fed with diets supplemented with 0.00015% MDP (MDP-1 group), 0.0015% MDP (MDP-2 group), and 0.015% MDP (MDP-3 group) for 7 days (6 replicate bottles for each group, 30 fish per bottle), respectively. (A-G) The effects of supplementing MDP to a diet on the growth performance of zebrafish. (A) Final body weight (FBW, n = 5). (B) Body length (n = 6). (C) Condition factor (CF, n = 6). (D) Viscerosomatic index (VSI, n = 6). (E) Carcass ratio (CR, n = 6). (F) Muscle moisture (n = 6). (G) Muscle protein content (n = 6). Data were presented as mean ± SEM. Student’s t-test was employed to compare between two groups. Significant differences were indicated as **p* < 0.05, ***p* < 0.01, ****p* < 0.001.

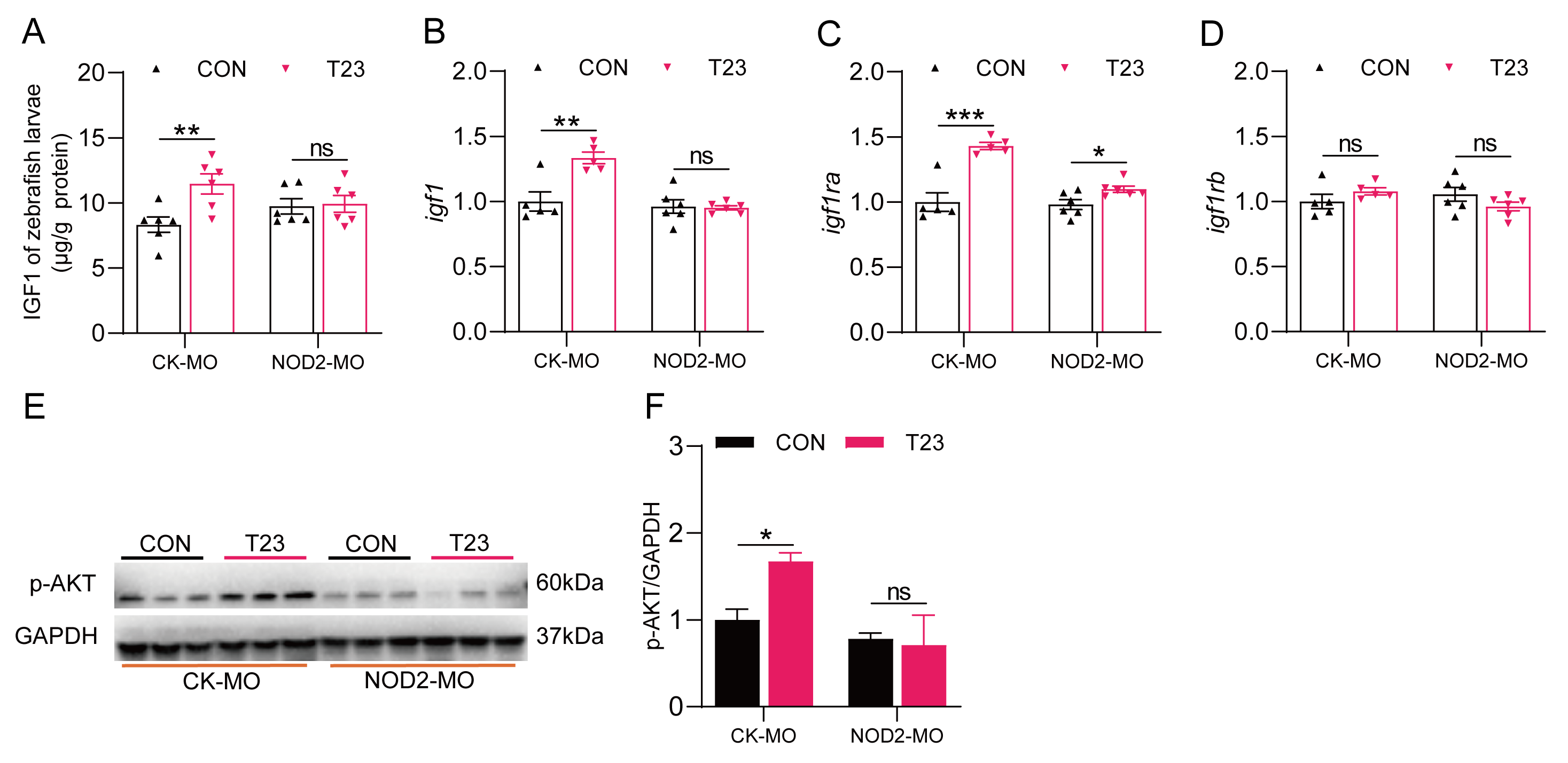

**Figure S6.** The effects of *B. velezensis* T23 on the IGF1 signaling pathway of zebrafish larvae treated with vivo morpholino directed to NOD2. Zebrafish larvae (CK-MO and NOD2-MO) were fed with diets supplemented with normal diet or diet supplemented with 10⁷ CFUs/g *B. velezensis* T23 for 7 days. (A) The content of IGF1 in the zebrafish larvae (n = 6). (B-D) The expression of growth-related genes (n = 5-6). (E) Western blot analysis and (F) quantitation of p-AKT in zebrafish larvae (n = 3). Data were presented as mean ± SEM. Student’s t-test was employed to compare between two groups. Significant differences were indicated as **p* < 0.05, ***p* < 0.01, ****p* < 0.001, ns=not significant.

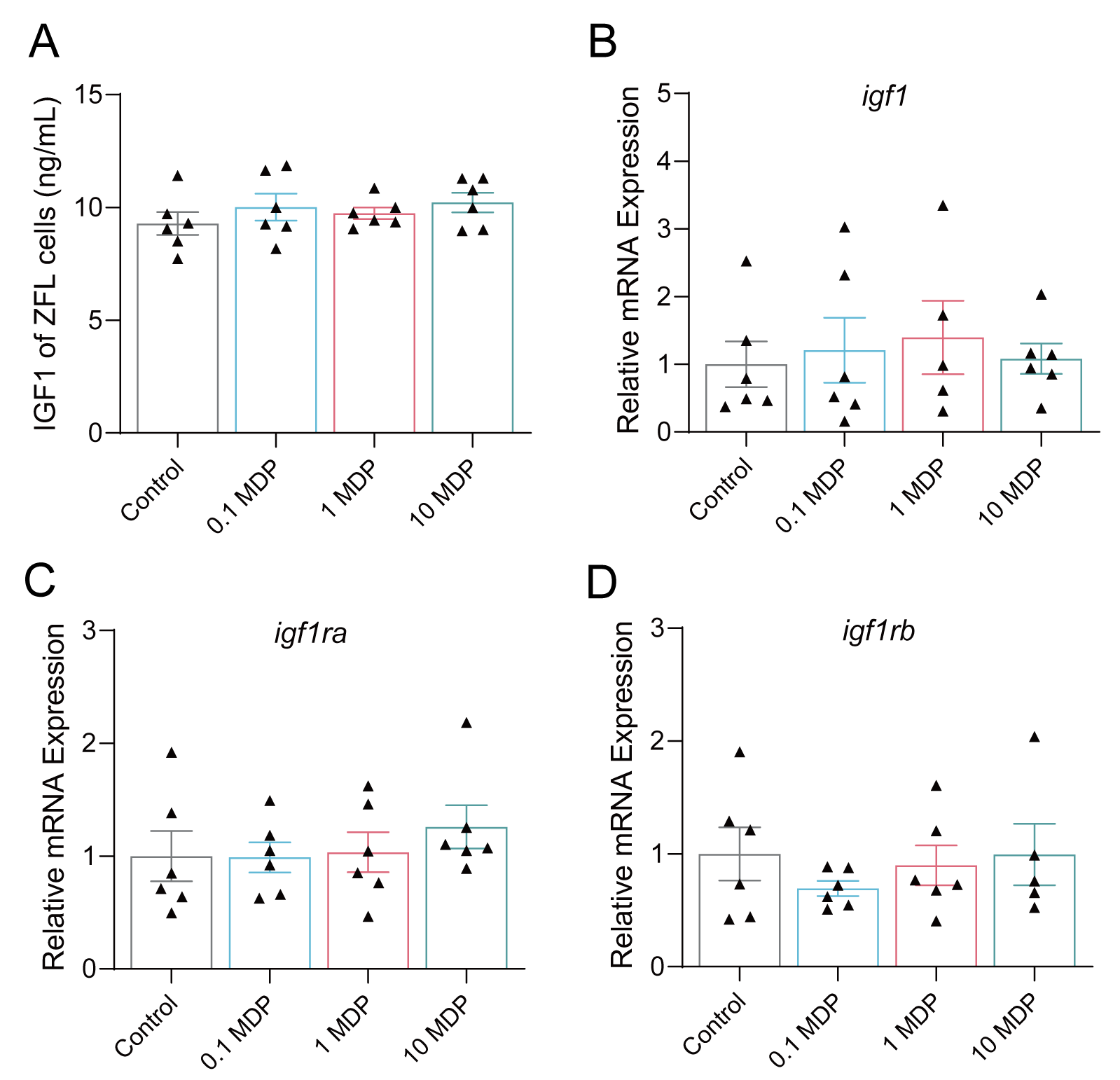

**Figure S7.** MDP was not able to directly activate the IGF1 signaling pathway in ZFL cells. ZFL cells were treated with MDP at final concentrations of 0.1, 1 and 10 μg/mL. (A) The content of IGF1 in ZFL cells (n = 6). (B-D) The expression of growth-related genes in ZFL cells (n = 6). Data were presented as mean ± SEM. Student’s t-test was employed to compare between two groups.

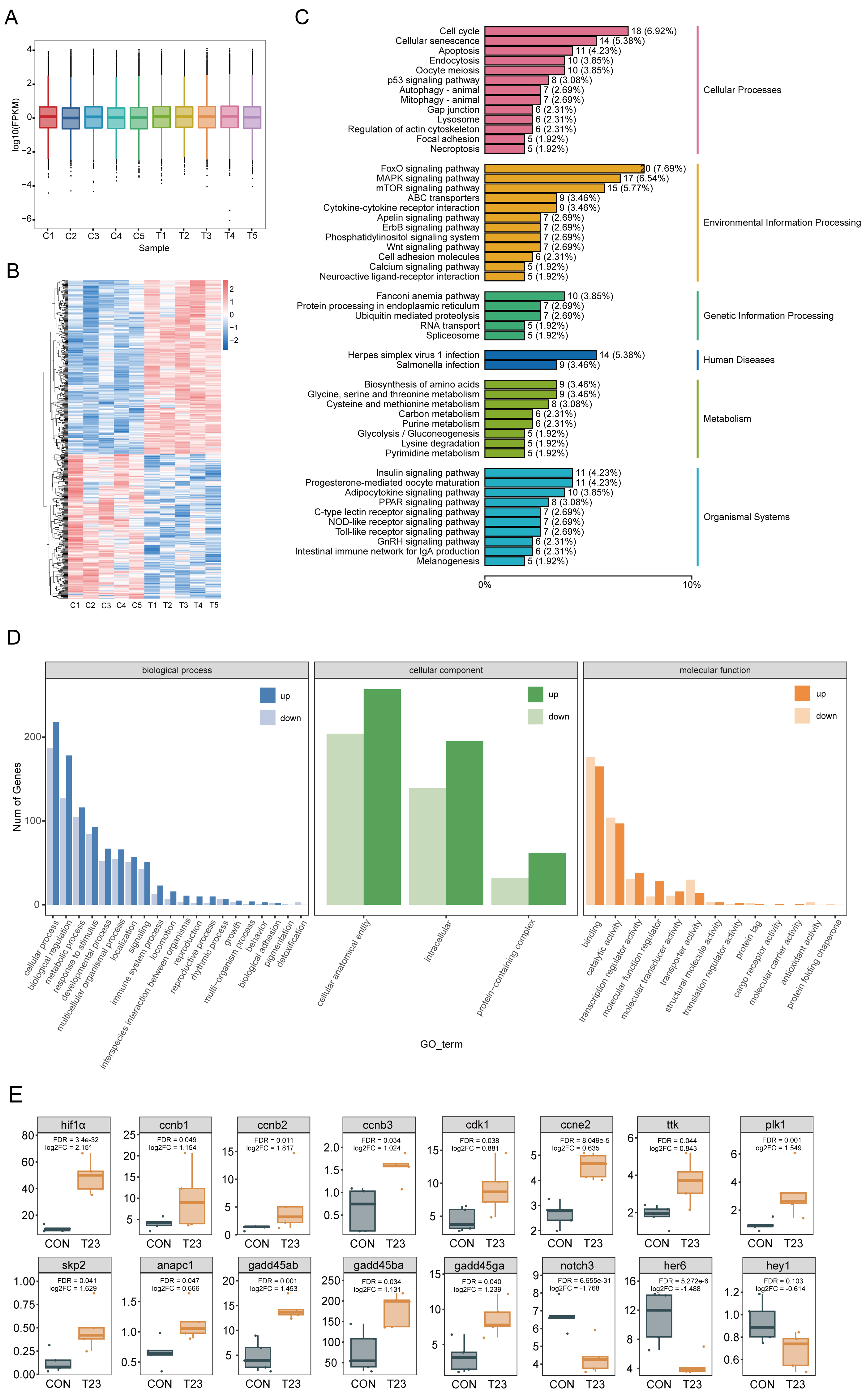

**Figure S8.** Transcriptome sequencing of the midgut of zebrafish after feeding with *B. velezensis* T23. (A) FPKM values of different samples (n = 5). (B) The heatmap showed the distribution of each sample for differentially expressed genes (DEGs) (Fold Change ≥ 1.5, FDR < 0.05). (D) KEGG classification annotation of DEGs. (D) GO enrichment analysis of DEGs. (E) FPKM values of the key differentially expressed genes in the two groups.

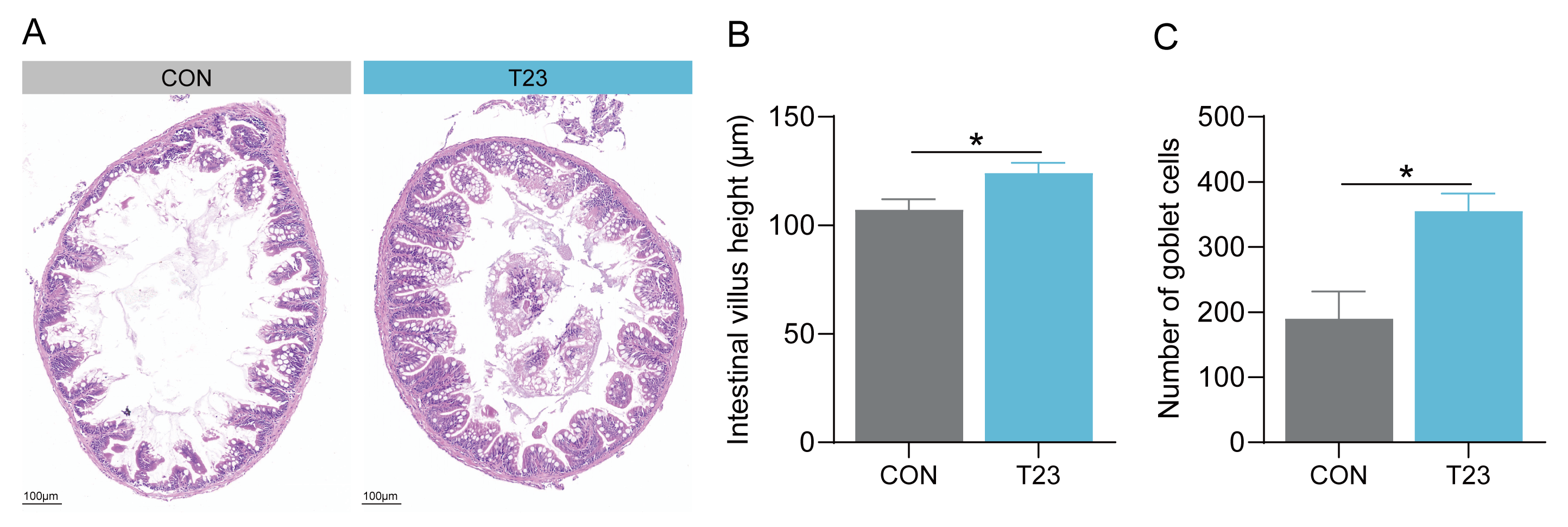

**Figure S9.** The effects of *B. velezensis* T23 on gut health of zebrafish. (A) Gut H&E section (n = 3). (B) Intestinal villus height (n = 3). (C) Number of goblet cells (n = 3). Data were presented as mean ± SEM. Student’s t-test was employed to compare between two groups. Significant differences were indicated as ***p* < 0.01.

**Supplemental Tables**

**Table S1.** Ingredients (g/100g diet) and chemical compositions (%) of zebrafish diets

| Ingredients (g/100g diet) | CON | T23 | T23-Δsfp | WCW | PGN | MDP |
| --- | --- | --- | --- | --- | --- | --- |
| Casein | 40.00 | 40.00 | 40.00 | 40.00 | 40.00 | 40.00 |
| Gelatin | 10.00 | 10.00 | 10.00 | 10.00 | 10.00 | 10.00 |
| Dextrin | 28.00 | 28.00 | 28.00 | 28.00 | 28.00 | 28.00 |
| Lard oil | 3.00 | 3.00 | 3.00 | 3.00 | 3.00 | 3.00 |
| Soybean oil | 3.00 | 3.00 | 3.00 | 3.00 | 3.00 | 3.00 |
| Lysine | 0.33 | 0.33 | 0.33 | 0.33 | 0.33 | 0.33 |
| VC phosphate | 0.10 | 0.10 | 0.10 | 0.10 | 0.10 | 0.10 |
| Vitamin premix^1^ | 0.40 | 0.40 | 0.40 | 0.40 | 0.40 | 0.40 |
| Mineral premix^2^ | 0.40 | 0.40 | 0.40 | 0.40 | 0.40 | 0.40 |
| Monocalcium phosphate | 2.00 | 2.00 | 2.00 | 2.00 | 2.00 | 2.00 |
| Choline chloride | 0.20 | 0.20 | 0.20 | 0.20 | 0.20 | 0.20 |
| Sodium alginate | 2.00 | 2.00 | 2.00 | 2.00 | 2.00 | 2.00 |
| Microcrystalline cellulose | 4.00 | 4.00 | 4.00 | 4.00 | 4.00 | 4.00 |
| Zeolite | 6.57 | 6.57 | 6.57 | 6.57 | 6.57 | 6.56985 |
| *B. velezensis* T23^3^ |  | 107 CFUs/g |  |  |  |  |
| *B. velezensis* T23-Δ*sfp*^4^ |  |  | 107 CFUs/g |  |  |  |
| Whole cell wall^5^ |  |  |  | Equal amount |  |  |
| Peptidoglycan^6^ |  |  |  |  | Equal amount |  |
| Muramyl dipeptide^7^ |  |  |  |  |  | 0.00015 |
| Total | 100.00 | 100.00 | 100.00 | 100.00 | 100.00 | 100.00 |
| Proximate analysis |  |  |  |  |  |  |
| Crude protein (%) | 42.19 | 42.19 | 42.19 | 42.19 | 42.19 | 42.19 |
| Crude fat (%) | 6.00 | 6.00 | 6.00 | 6.00 | 6.00 | 6.00 |

1. Containing the following (g/kg vitamin premix): thiamine, 0.438; riboflavin, 0.632; pyridoxine⋅HCl, 0.908; d-pantothenic acid, 1.724; nicotinic acid, 4.583; biotin, 0.211; folic acid, 0.549; vitamin B-12, 0.001; inositol, 21.053; menadione sodium bisulfite, 0.889; retinyl acetate, 0.677; cholecalciferol, 0.116; dl-α-tocopherol-acetate,12.632.

2. Containing the following (g/kg mineral premix): CoCl_2_⋅6H_2_O, 0.074; CuSO_4_⋅5H_2_O, 2.5; FeSO_4_⋅7H_2_O, 73.2; NaCl, 40.0; MgSO_4_⋅7H_2_O, 284.0; MnSO_4_⋅H_2_O, 6.50; KI, 0.68; Na_2_SeO_3_, 0.10; ZnSO_4_⋅7H_2_O, 131.93; Cellulose, 501.09.

3. *Bacillus velezensis* T23 (*B. velezensis* T23).

4. *Bacillus velezensis* T23-Δ*sfp* (*B. velezensis* T23-Δ*sfp*).

5. Whole cell wall was extracted from an equal amount of *B. velezensis* T23.

6. Peptidoglycan was extracted from an equal amount of *B. velezensis* T23.

7. Muramyl dipeptide was purchased from MCE, USA.

**Table S2.** Ingredients (g/100g diet) and chemical compositions (%) of zebrafish larvae diets

| Ingredients (g/100g diet) | CON | T23 |
| --- | --- | --- |
| Casein | 46.00 | 46.00 |
| Gelatin | 11.00 | 11.00 |
| Dextrin | 22.00 | 22.00 |
| Soybean oil | 3.50 | 3.50 |
| Fish liver oil | 3.50 | 3.50 |
| Soybean lecithin | 2.00 | 2.00 |
| Lysine | 0.37 | 0.37 |
| VC phosphate | 0.10 | 0.10 |
| Vitamin premix^1^ | 0.40 | 0.40 |
| Mineral premix^2^ | 0.40 | 0.40 |
| Monocalcium phosphate | 2.00 | 2.00 |
| Choline chloride | 0.20 | 0.20 |
| Sodium alginate | 4.00 | 4.00 |
| Zeolite | 4.53 | 4.53 |
| *B. velezensis* T23^3^ |  | 10^7^ CFUs/g |
| Total | 100.00 | 100.00 |
| Proximate analysis |  |  |
| Crude protein (%) | 48.09 | 48.09 |
| Crude fat (%) | 22.00 | 22.00 |

1. Containing the following (g/kg vitamin premix): thiamine, 0.438; riboflavin, 0.632; pyridoxine⋅HCl, 0.908; d-pantothenic acid, 1.724; nicotinic acid, 4.583; biotin, 0.211; folic acid, 0.549; vitamin B-12, 0.001; inositol, 21.053; menadione sodium bisulfite, 0.889; retinyl acetate, 0.677; cholecalciferol, 0.116; dl-α-tocopherol-acetate,12.632.

2. Containing the following (g/kg mineral premix): CoCl_2_⋅6H_2_O, 0.074; CuSO_4_⋅5H_2_O, 2.5; FeSO_4_⋅7H_2_O, 73.2; NaCl, 40.0; MgSO_4_⋅7H_2_O, 284.0; MnSO_4_⋅H_2_O, 6.50; KI, 0.68; Na_2_SeO_3_, 0.10; ZnSO_4_⋅7H_2_O, 131.93; Cellulose, 501.09.

3. *Bacillus velezensis* T23 (*B. velezensis* T23).

**Table S3.** List of primer sequences used for qPCR

| Primer | Forward primer (5′ –3′) | Reverse primer (5′ –3′) | Reference |
| --- | --- | --- | --- |
| *rps11* | ACAGAAATGCCCCTTCACTG | GCCTCTTCTCAAAACGGTTG | [1] |
| *igf1* | CCTAGTTCAAGAAGGTCACACAAC | CTATGCCCAGATATAGGTTTCTTTGGT | This study |
| *igf1ra* | GCCCGTGGAGAAGTCTGTGG | GTGTGCGAAAGTGTTCCTGGTT | [2] |
| *igf1rb* | ATCCTCCCGGCCTTACTGTT | CCTGTCATTGTTTCGGTTCTTGT | [2] |
| *ghra* | GGCCGAAAATTCCTTACTGTT | GCTGGCGTTGCTGATTGT | [2] |
| *ghrb* | GCTGCGCTCTGTTGATAATGT | GGCGGAGGGAGGTGGAT | [2] |
| *B. velezensis* | GTCCGGGGGCATTGGCTGAG | CCCCCTGCACATAGACGGACTGA | [3] |

*rps11*, ribosomal protein S11; *igf1*, insulin-like growth factor 1; *igf1ra*, insulin-like growth factor 1a; *igf1rb*, insulin-like growth factor 1b; *ghra*, growth hormone receptor a; *ghrb*, growth hormone receptor b.
